# Intermediate phenotypic heterogeneity creates an optimal coexistence window in bacteria–phage communities

**DOI:** 10.64898/2026.10.01.755942

**Authors:** Jung-Ho Kim, Mattia Mattei, David Soriano-Paños, Alex Arenas

## Abstract

Phenotypic heterogeneity can expand ecological niche space even within a genetically homogeneous population. Recent experiments showed that growth-rate heterogeneity in a clonal bacterial host can stabilize coexistence among multiple phages. Yet this discovery leaves unresolved whether increasing host heterogeneity monotonically promotes coexistence. Here we show that it does not. Using a trait-based bacteria–phage coexistence framework, we find that phage richness is maximized at an intermediate scale of host growth-rate heterogeneity, determined by the match between host phenotypic dispersion and phage niche width. Our results show that optimal phage diversity emerges from an ecological trade-off between two competing effects: increasing phenotypic heterogeneity enables niche differentiation but eventually reduces the host abundance in the preferred niche of each phage. Therefore, our findings predict that experimentally increasing host growth-rate variance at a fixed mean growth rate should produce a nonmonotonic phage richness response, providing a quantitative principle for how intraspecific phenotypic variation shapes biodiversity in microbial ecosystems.

**Significance statement:** Genetically homogeneous bacterial populations can contain phenotypically distinct subpopulations, creating different ecological opportunities for the bacteriophages that infect and prey on them. However, we show that this heterogeneity does not simply increase phage diversity. Instead, phage diversity is greatest at an intermediate level of bacterial phenotypic heterogeneity. Too little heterogeneity leaves phages competing for similar bacteria, whereas too much spreads bacterial biomass across phenotypic space, leaving phages with too few suitable hosts to persist. The optimum arises when host phenotypic dispersion matches phage infection breadth. This scale-matching principle links variation within a single population to biodiversity among interacting species and provides a direct experimental prediction for how microbial communities should respond to changes in phenotypic heterogeneity.

---

Understanding how bacteriophages exploit microbial populations to generate and maintain high levels of diversity in natural ecosystems is a central question in theoretical ecology (*1–4*). In well-mixed environments, classical consumer–resource theory predicts that the number of coexisting phage strains cannot exceed the number of distinct bacterial strains (*5–7*). This upper bound is attained when the ecological niches (*8, 9*) of the surviving phages—that is, the bacterial hosts they infect—do not overlap, thereby minimizing competition for limited resources. Guided by this principle, considerable theoretical effort has focused on identifying mechanisms that enhance niche differentiation, revealing that spatial structure (*10–13*), temporal fluctuations in resource availability (*14, 15*), and bacterial diversification (*16–18*) can all promote phage diversity.

According to the consumer-resource model, a clonal bacterial population should not, by itself, provide the ecological niche structure required to sustain multiple competing phages. Yet recent experiments showed that multiple phage species can coexist on a genetically homogeneous bacterial host because phenotypic growth-rate heterogeneity creates effective ecological niches which can be exploited by different phages (*19*). This finding raises a sharper theoretical question: does increasing host phenotypic heterogeneity always promote phage diversity (*20–22*)?

To address this question, we develop a generalized Lotka–Volterra framework (*17, 23–28*) in which bacterial phenotypes differ in their intrinsic growth rates and phage infection efficiency depends on the mismatch between the host growth rate and their preferred niche. We first validate our framework, showing that bacterial phenotypic heterogeneity and a trait-based interaction kernel allow reproducing both the phage diversity and the host-state-dependent turnover in phage abundance observed in real experiments (*19*).

By extending this framework to generalized bacteria–phage ensembles, we investigate how phage coexistence depends on bacterial phenotypic heterogeneity and phage niche width. We further quantify the underlying coexistence constraints through niche differentiation (ND) and invasion viability (IV), which measures the fraction of host–phage pairs with sufficient infection-weighted host biomass for invasion. This framework allows us to determine how the relative scales of host phenotypic dispersion and phage niche width shape phage coexistence.

## Generalized Lotka–Volterra dynamics

To formulate the temporal dynamics of our model, we consider a well-mixed population of bacteria and phages and employ a generalized Lotka–Volterra framework (*17, 23–28*) [Fig. 1(a)] incorporating: *i*) Logistic growth of each bacterial phenotype *i* governed by its intrinsic growth rate *r*_*i*_ and the carrying capacity *C, ii*) dilution of the bacterial population at a rate *v, iii*) phage infection processes governed by the affinity matrix **A** and the phage burst size *ξ* and *iv*) phage degradation at a rate *μ*.

**Figure 1:**
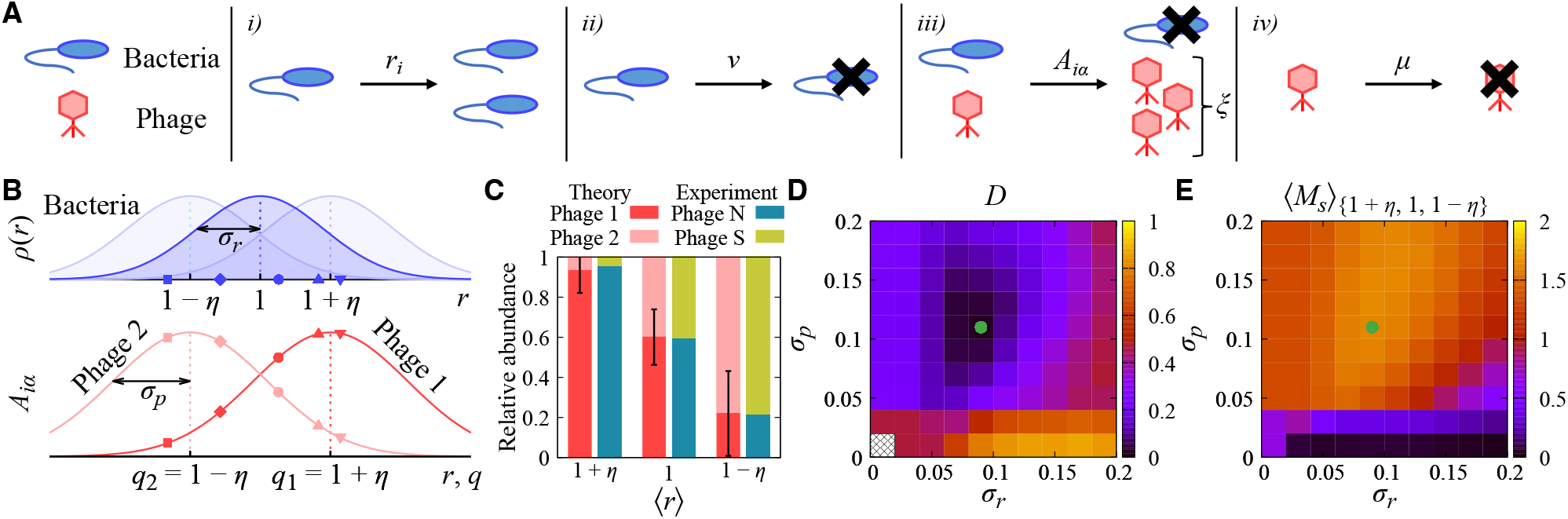
Experimental grounding of the minimal growth-rate-dependent framework. (**A**) Schematic diagram of the generalized Lotka–Volterra dynamics. (**B**) Two-phage setup used to compare the model with the experimentally observed abundance turnover. (Top panel) The three shaded distributions represent host populations with high, intermediate, and low mean growth rates, ⟨*r*⟩ = 1 + *η*, 1, and 1 − *η*, respectively. Marked bacterial phenotypes indicate one representative realization sampled from the intermediate-growth population with ⟨*r*⟩ = 1. (Bottom panel) The corresponding affinity values *A*_*iα*_ are shown for Phage 1, with preferred host growth rate *q*_1_ = 1 +*η*, and Phage 2, with preferred host growth rate *q*_2_ = 1 − *η*. (**C**) Relative abundance turnover across the three host growth states. The theoretical bars show the relative abundances obtained from the minimal two-phage framework with *η* = 0.1, *σ*_*r*_ = 0.09, and *σ*_*p*_ = 0.11. The experimental bars show the corresponding relative abundances adapted from Pyenson et al. (*19*). Error bars denote the standard deviations. (**D**) Euclidean distance *D* between the experimental relative-abundance vector across the high, intermediate, and low growth states and the corresponding model-predicted vector across ⟨*r*⟩ = 1 + *η*, 1, and 1 − *η*, assuming *η* = 0.1. White cross-hatched cells indicate parameter sets for which no phage survives at ⟨*r*⟩ = 1; the phage relative abundance, and consequently *D*, is therefore undefined. The green dot highlights the parameter set used in **C**. (**E**) Average number of surviving phage species ⟨*M*_*s*_⟩_{1+*η*, 1, 1−*η*}_ in the same (*σ*_*r*_, *σ*_*p*_) parameter space. For each parameter set, ⟨*M*_*s*_⟩_{1+*η*, 1, 1−*η*}_ is averaged over the three host-growth regimes, ⟨*r*⟩ = 1 + *η*, 1, and 1 − *η*. Results are averaged over 2 × 10^3^ independent realizations.

We treat each bacterial phenotypic state as a persistent subpopulation over the ecological timescale described by the model. Accordingly, we use a coarse-grained ecological assumption that bacterial growth within phenotype *i* contributes to the same phenotypic class. This approximation is consistent with the experimental setting motivating our model, in which phage coexistence was attributed primarily to host heterogeneity already present at the onset of each passage rather than to phenotypic changes generated during the subsequent bacteria–phage dynamics (*19*).

To compute the affinity matrix, we assume that bacterial growth rates determine the ecological niches of phages. Specifically, we set a host growth rate *q*_*α*_ preferred by each phage *α* and assume a Gaussian infection kernel (*16, 29–31*) of width *σ*_*p*_ around that value. Consequently, the infection efficiency peaks when the bacterial growth rate matches the phage preference (*r*_*i*_ = *q*_*α*_) and decays symmetrically with phenotypic mismatch. The equations governing the time evolution of the system and the parameter values used in the simulations are given in Materials and Methods, with additional numerical details provided in the Supplementary Information.

### Experimental grounding of the minimal framework

We first examine whether the minimal growth-rate-dependent interaction kernel can recapitulate the empirical results reported in (*19*). There, the relative abundances of two coexisting phages are reversed when these phages infect bacterial populations characterized by high, intermediate, and low average growth states. Motivated by this setup, we perform three *in silico* experiments, assuming three bacterial populations with different average growth rates: ⟨*r*⟩ = 1 + *η*, 1, or 1 − *η*, representing high, intermediate, and low growth states, respectively [Fig. 1(b)]. In each experiment, we sample *N* = 10 bacterial phenotypes from a Gaussian distribution with phenotypic heterogeneity *σ*_*r*_ and infect them with two phage species whose optimal host growth rates are set to *q*_1_ = 1 + *η* and *q*_2_ = 1 − *η*, corresponding to specialists on fast- and slow-growing host phenotypes, respectively.

Despite its simplicity, our *in silico* experiment recapitulates the reversal in relative phage abundance as the host population shifts from fast- to slow-growth regimes reported in the empirical experiments [Fig. 1(c)]. Therefore, a simple growth-rate-dependent infection kernel is sufficient to generate the observed host-state-dependent abundance turnover.

We next study how both this agreement with the experiment and the phage diversity found in the *in silico* model depend on bacterial phenotypic heterogeneity, dictated by *σ*_*r*_, and the phage niche specialization, dictated by *σ*_*p*_. We find a broad region of the parameter space compatible with the abundance turnover reported empirically [Fig. 1(d)]. Importantly, this experimentally grounded region also shows enhanced phage coexistence, as indicated by the elevated average number of surviving phage species, ⟨*M*_*s*_⟩_{1+*η*, 1, 1−*η*}_, obtained across the three host-growth regimes [Fig. 1(e)]. Nevertheless, when *σ*_*r*_ ≫ *σ*_*p*_, i.e. for highly heterogeneous bacterial populations, we find that our simulations no longer reproduce the empirical abundance turnover and predict a reduction in the number of surviving phages, suggesting that excessive phenotypic diversity can hamper phage diversity. To shed light on this effect, we next consider generalized bacteria–phage ensembles where we can systematically investigate how host phenotypic diversity and phage specificity shape phage diversity.

### Phage diversity in generalized bacteria–phage ensembles

We next extended the framework to generalized bacteria–phage ensembles to examine how phage diversity depends on host phenotypic dispersion and phage niche width. In this ensemble setting, both the bacterial growth rates *r*_*i*_ (*i* = 1, …, *N* = 10) and the phage preferred host states *q*_*α*_ (*α* = 1, …, *M* = 10) were treated as random variables drawn from Gaussian trait distributions. Because host and phage traits are expected to occupy comparable ranges after long-term ecological or evolutionary matching, we set ⟨*r*⟩ = ⟨*q*⟩ = 1 and *σ*_*r*_ = *σ*_*q*_ = *σ*, so that *σ* controls the common scale of host and phage trait dispersion [Fig. 2(a)].

**Figure 2:**
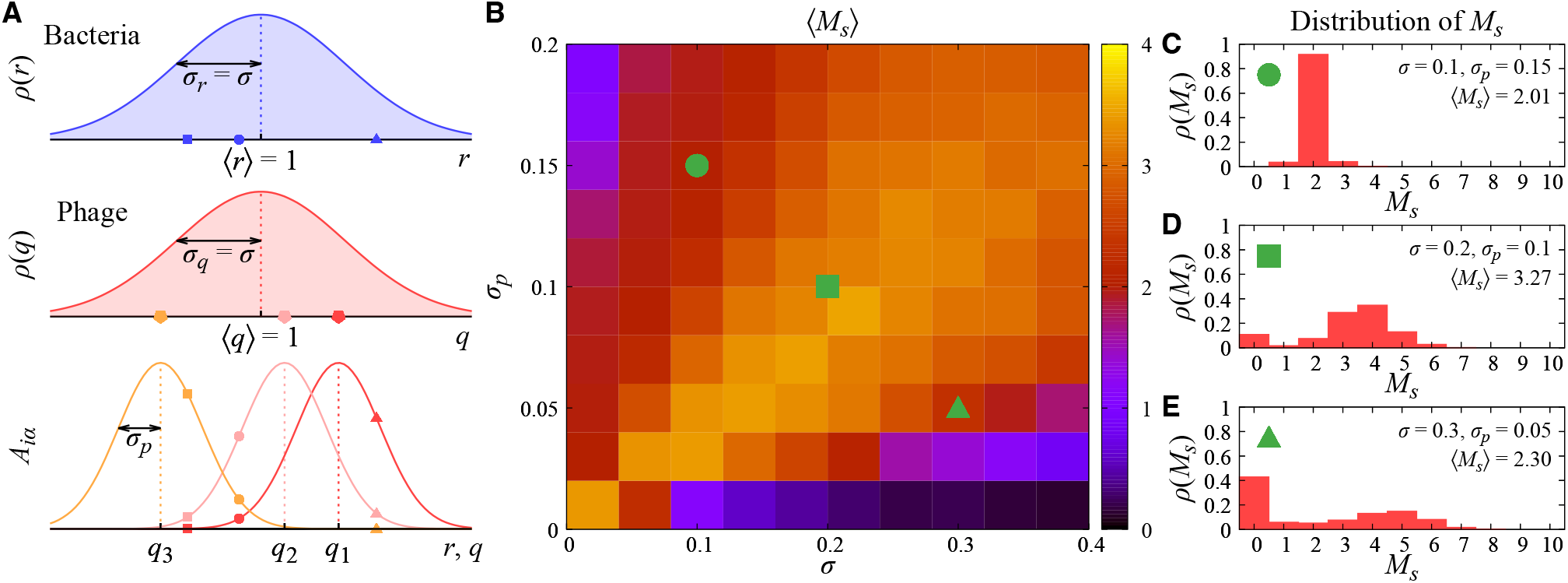
Phage diversity in generalized bacteria–phage ensembles. (**A**) Schematic illustration of the generalized bacteria-phages ensemble. Bacterial growth rates {*r*} and phage preferred host states {*q*} are independently sampled from Gaussian trait distributions with matched mean and variance, ⟨*r*⟩ = ⟨*q*⟩ = 1 and *σ*_*r*_ = *σ*_*q*_ = *σ*. The infection efficiency *A*_*iα*_ of phage *α* for bacteria *i* is determined by the growth-rate-dependent infection kernel with phage niche width *σ*_*p*_. (**B**) Average number of surviving phage species ⟨*M*_*s*_⟩, as a function of host/phage trait dispersion *σ* and phage niche width *σ*_*p*_. (**C**–**E**) Distributions of the number of surviving phage species *M*_*s*_ for representative parameter sets indicated in (**B**): (**C**) low trait dispersion and broad phage niche width, *σ* = 0.1 and *σ*_*p*_ = 0.15; (**D**) intermediate scale matching, *σ* = 0.2 and *σ*_*p*_ = 0.1; and (**E**) high trait dispersion and narrow phage niche width, *σ* = 0.3 and *σ*_*p*_ = 0.05. Results are averaged over 10^3^ independent realizations.

Phage diversity ⟨*M*_*s*_⟩, understood as the average number of surviving phages at long times across multiple realizations, shows a nonmonotonic behavior both with host phenotypic dispersion *σ* and phage specificity *σ*_*p*_ [Fig. 2(b)]. Namely, diversity is maximized when both scales match, i.e. *σ* ∼ *σ*_*p*_ whereas fewer phages survive when phenotypic diversity is much smaller or much greater than the phage niche width, i.e. when either *σ* ≪ *σ*_*p*_ or *σ* ≫ *σ*_*p*_ respectively. Phage coexistence capacity is low when host phenotypic dispersion is small and phage niches are broad, and it is also low when host dispersion is large while phage niches are narrow. This qualitative pattern is robust to community size, model details, and the assumption of phenotypic persistence: the same nonmonotonic trend is observed across alternative infection kernels and trait distributions, different community sizes, and an opposite limiting model in which offspring phenotypes are randomly reassigned at reproduction as shown in Supplementary Text.

The full distribution of *M*_*s*_ further shows that the two low-diversity regimes are not equivalent [Figs. 2(c)–2(e)]. When *σ* is small and *σ*_*p*_ is large, the distribution of *M*_*s*_ remains relatively narrow, indicating that most realizations produce small but typically nonzero numbers of surviving phages. By contrast, when *σ* is large and *σ*_*p*_ is small, the distribution broadens and includes many realizations with zero surviving phages. These distributional differences motivate a direct quantification of the two opposing constraints that shape the coexistence window: whether phage niches are sufficiently differentiated, and whether enough viable host–phage links remain to sustain infection.

### Mechanistic origin of the optimal coexistence window

To identify the mechanism underlying the optimal coexistence window, we quantified two opposing constraints on phage coexistence: niche differentiation (ND) and invasion viability (IV). ND measures how strongly phage infection kernels are separated along the host growth-rate axis. Thus, small ND indicates that phage niches strongly overlap, whereas large ND indicates that phages target distinct regions of phenotypic space. By contrast, IV measures the fraction of host–phage pairs that are individually viable for invasion. A host–phage pair is counted as viable when the infection-weighted host biomass available to the phage exceeds the phage maintenance threshold, evaluated in the corresponding pairwise invasion setting. Formal definitions of ND and IV are provided in Materials and Methods, with analytical expressions given in the Supplementary Information.

Mapping both quantities across the same (*σ, σ*_*p*_) parameter space reveals the two sides of the coexistence trade-off [Figs. 3(a) and 3(b)]. When host phenotypic dispersion is much smaller than phage niche width (*σ* ≪ *σ*_*p*_), IV is high but ND remains low, leading to strong competition among phages. This explains the low phage diversity in Fig. 2(b), as well as the relatively narrow *M*_*s*_ distribution in Fig. 2(c), where realizations with *M*_*s*_ = 0 are rare. Conversely, when host phenotypic dispersion is large and phage niche width is narrow, i.e. *σ* ≫ *σ*_*p*_, ND is high but IV declines sharply. In this regime, phages are highly specialized, but many fail to find host phenotypes that provide enough infection-weighted biomass to maintain growth. This accounts for the broad *M*_*s*_ distribution with many zero-survival realizations observed in Fig. 2(e). The optimal coexistence window therefore emerges between these limits, where phage niches are sufficiently differentiated while enough viable host–phage links remain to sustain infection.

**Figure 3:**
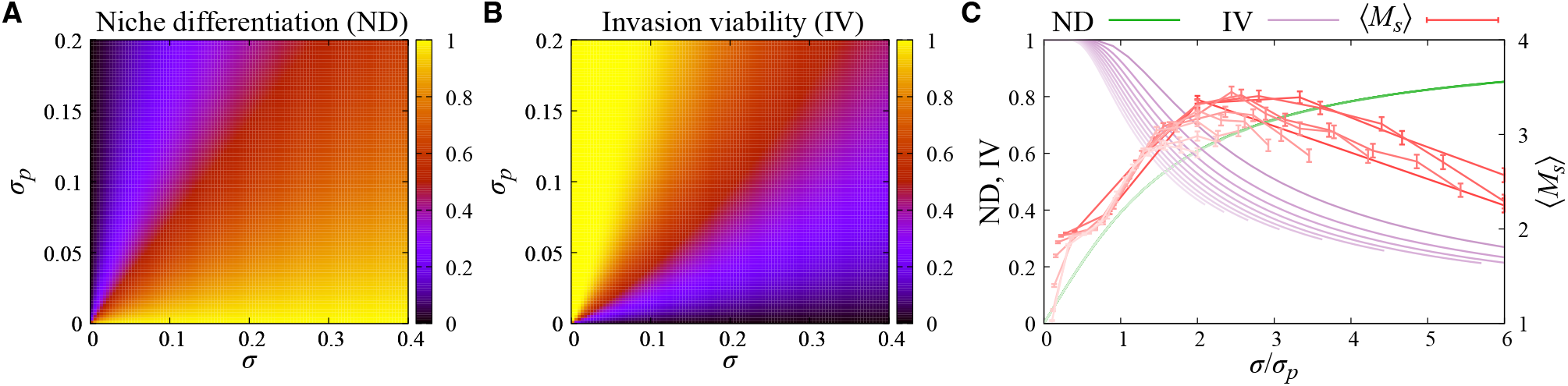
Niche differentiation and invasion viability explain the optimal coexistence window. (**A**) Niche differentiation (ND) as a function of the phenotypic heterogeneity *σ* and the phage niche width *σ*_*p*_. (**B**) Invasion viability (IV) in the same parameter space. (**C**) ND, IV, and coexistence capacity ⟨*M*_*s*_⟩ plotted as functions of the scale ratio *σ*/*σ*_*p*_. Each curve corresponds to a fixed value of *σ*_*p*_ from 0.01 to 0.19 in increments of 0.02, with darker curves indicating smaller *σ*_*p*_. Error bars for ⟨*M*_*s*_⟩ denote the standard errors.

The scale-matching nature of this mechanism becomes clearer when ND, IV, and ⟨*M*_*s*_⟩ are plotted as functions of the dimensionless ratio *σ*/*σ*_*p*_ [Fig. 3(c)]. Regardless of the *σ*_*p*_ value, phage diversity is optimized when both scales are comparable, as both ND and IV take intermediate values that promote the survival of phages. Together, these patterns support a scale-matching principle: phage coexistence is optimized when host phenotypic dispersion is neither too small nor too large relative to phage infection breadth.

Finally, to confirm the role of this ecological trade-off in maximizing phage diversity, we analytically solve a minimal two-host–two-phage model in the Supplementary Text. Our analytical results show that the optimal coexistence window also emerges in this simple communities, thus ruling out stochastic sampling of both bacterial phenotypes and phage species as the main driver of the phenomenology observed in general bacteria-phage ensembles.

## Conclusion

Phenotypic heterogeneity is a fundamental determinant of the resilience and adaptability of clonal bacterial populations in fluctuating environments (*32–34*). Beyond conferring ecological advantages to bacteria, recent experimental evidence from bacteria–phage communities has shown that such heterogeneity can also create ecological niches that support the coexistence of multiple phage species (*19*). Motivated by these findings, we developed and analyzed a minimal theoretical framework to elucidate how bacterial phenotypic heterogeneity, together with phage specialization, shapes the emergence and maintenance of phage diversity.

We first validated our framework by reproducing in silico the experiments reported in (*19*), obtaining good quantitative agreement between our theoretical predictions and the empirical observations. Combining simulations of general bacteria–phage communities with analytical results for simplified systems, we found that phage diversity is maximized at intermediate levels of bacterial phenotypic heterogeneity. This optimal coexistence regime emerges from an ecological trade-off. At low levels of phenotypic heterogeneity, the host population is insufficiently differentiated to generate distinct ecological niches for competing phages. Increasing heterogeneity promotes niche partitioning and thus coexistence, but once heterogeneity becomes sufficiently large, the effective host biomass available to each specialized phage becomes too small to sustain successful invasion.

Our findings provide a mechanistic principle for how microscopic, intraspecific variation shapes macroscopic microbial biodiversity. Our results also yield direct experimental predictions: increasing host growth-rate variance at a fixed mean growth rate should generate a nonmonotonic phage richness response, and the location of the optimum should shift with phage infection breadth. An important challenge for future empirical and theoretical studies will be to determine whether real bacteria–phage communities operate within this optimal coexistence regime and, if so, whether this regime emerges as a consequence of the coevolutionary dynamics between hosts and their phages (*35–37*).

While the current model focuses on a well-mixed environment with a fixed trait distribution, future extensions incorporating spatial heterogeneity (*10–13*), explicit bacteria–phage interaction networks (*16,26,27,38–42*), dynamic trait evolution (*16,30,31,43–45*), or experimentally measured phenotypic distributions (*46*) will further illuminate the complex realities of natural host–virus interactions.

Ultimately, this work highlights that phenotypic heterogeneity is not merely biological noise, but a critical structural determinant whose scale controls microbial coexistence.

## Materials and Methods

### Generalized Lotka–Volterra dynamics

We modeled a well-mixed community of bacterial phenotypes and phage species using a generalized Lotka–Volterra framework (*17, 23–28*). Let *B*_*i*_ denote the density of bacterial phenotype *i* (*i* = 1, …, *N*), and let *P*_*α*_ denote the density of phage species *α* (*α* = 1, …, *M*). The dynamics are given by

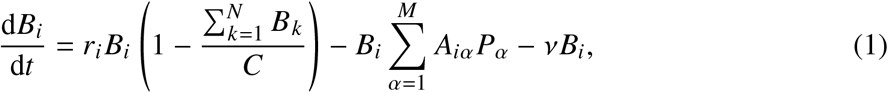

and

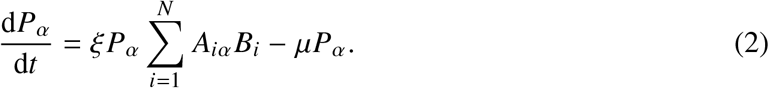

Here, *r*_*i*_ is the intrinsic growth rate of bacterial phenotype *i, C* is the bacterial carrying capacity, *v* is the bacterial dilution or mortality rate, *ξ* is the phage burst size, and *μ* is the phage degradation rate. The affinity matrix **A** encodes growth-rate-dependent infection efficiency. In the main analysis, we used the Gaussian infection kernel defined as

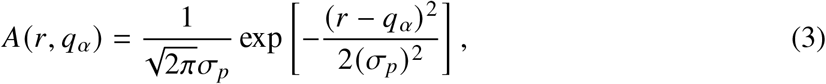

where each phage *α* preferentially infects bacterial phenotypes with growth rates close to its preferred host state *q*_*α*_.

The dynamics were integrated numerically using a fourth-order Runge–Kutta method with time step Δ*t* = 0.01. Unless otherwise stated, we set *C* = 1, *v* = 0.1, *μ* = 1, and *ξ* = 2. All bacterial and phage densities were initialized as *B*_*i*_ (0) = *P*_*α*_ (0) = 0.01. The phage diversity reported in the main text was quantified by the ensemble-averaged number of surviving phage species, ⟨*M*_*s*_⟩, computed over independent random trait realizations at final integration times determined from convergence tests. Details of the numerical methods are provided in the Supplementary Information.

### Niche differentiation and invasion viability

To identify the mechanism underlying the optimal coexistence window, we quantified two complementary constraints on phage coexistence: niche differentiation (ND) and invasion viability (IV).

ND measures the average separation between pairs of phage infection kernels along the bacterial growth-rate axis. For two phages with preferred host states *q*_*α*_ and *q*_*β*_, we defined the pairwise niche differentiation as the normalized non-overlapping area between their infection kernels,

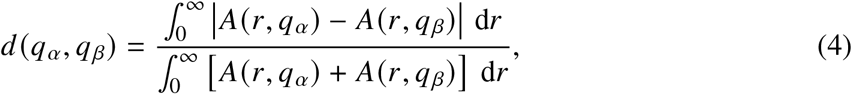

where *A*(*r, q*_*α*_) denotes the continuous infection kernel of phage *α* as a function of host growth rate *r*. The affinity matrix used in the dynamical equations is obtained by evaluating this kernel at the sampled bacterial growth rates, *A*_*iα*_ = *A*(*r*_*i*_, *q*_*α*_). This quantity ranges from zero for completely overlapping niches to one for non-overlapping niches.

The niche differentiation was then obtained by averaging this quantity over two independently sampled phage preferred host states,

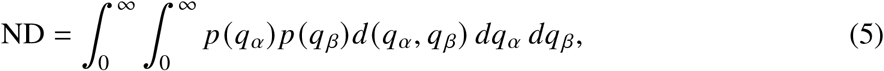

where *p* (*q*) denotes the phage-preference distribution.

IV measures the fraction of host–phage pairs that are individually viable for invasion. For a bacterial phenotype with growth rate *r*, the phage-free equilibrium density in the corresponding pairwise system is

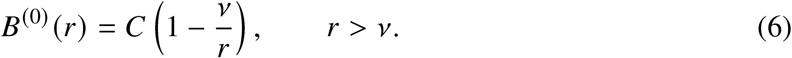

A rare phage with preferred host state *q*_*α*_ can invade this host phenotype when its initial per-capita growth rate is positive,

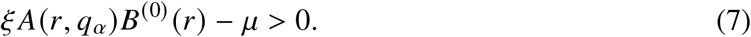

Thus, invasion viability was defined as

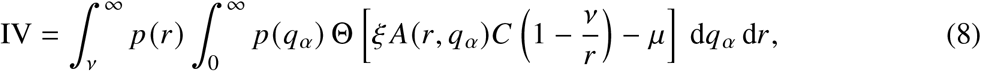

where *p* (*r*) and *p* (*q*_*α*_) denote the bacterial growth-rate and phage-preference distributions, respectively, and Θ(·) is the Heaviside step function.

ND therefore quantifies whether phage niches are sufficiently separated, whereas IV quantifies whether enough host–phage links remain viable for invasion. Analytical expressions for ND and IV for the Gaussian and log-normal infection kernels are provided in the Supplementary Information.

## Author contributions

Conceptualization: J.-H.K., M.M., D.S.-P., A.A., Data curation: J.-H.K., M.M., D.S.-P., A.A., Formal analysis: J.-H.K., M.M., D.S.-P., A.A., Funding acquisition: A.A., Investigation: J.-H.K., M.M., D.S.-P., A.A., Methodology: J.-H.K., M.M., D.S.-P., A.A., Project administration: A.A., Resources: A.A., Software: J.-H.K., Supervision: A.A., Validation: J.-H.K., M.M., D.S.-P., A.A., Visualization: J.-H.K., Writing – original draft: J.-H.K., Writing – review & editing: M.M., D.S.-P., A.A.

## Data, code and materials availability

The source codes can be freely accessed at the GitHub repository: (*47*).

## Supplementary Materials

### Supplementary Methods

#### Mathematical model for bacteria-phage dynamics

##### Model equations

We simulated well-mixed bacteria–phage communities using the generalized Lotka–Volterra dynamics described in the main text. Let *B*_*i*_ denote the density of bacterial phenotype *i* (*i* = 1, …, *N*), and let *P*_*α*_ denote the density of phage species *α* (*α* = 1, …, *M*). The carrying capacity *C* sets the scale of bacterial densities, and we set *C* = 1 in all numerical simulations unless otherwise stated. The dynamics are given by

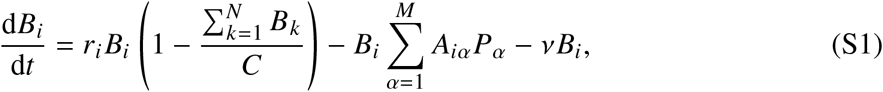

and

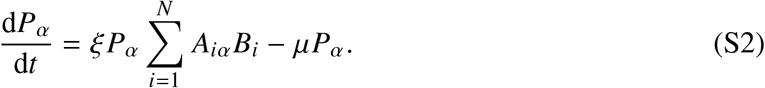

Here, *r*_*i*_ is the intrinsic growth rate of bacterial phenotype *i, v* is the bacterial dilution or mortality rate, *ξ* is the phage burst size, and *μ* is the phage degradation rate. The affinity matrix **A** has entries *A*_*iα*_ that quantify the infection efficiency between bacterial phenotype *i* and phage species *α*. The Gaussian trait distribution and infection kernel used in the main text to construct the bacterial growth rates, phage preferred host states, and affinity matrix are defined below.

##### Numerical methods

We numerically integrated Eqs. (S1) and (S2) using a fourth-order Runge– Kutta method with time step Δ*t* = 0.01. Unless otherwise stated, the demographic parameters were fixed to *v* = 0.1, *μ* = 1, and *ξ* = 2. For each random trait realization, all bacterial and phage densities were initialized as *B*_*i*_ (0) = 0.01 and *P*_*α*_ (0) = 0.01, for all bacterial phenotypes *i* and phage species *α*. During numerical integration, any bacterial phenotype or phage species whose density fell below the extinction threshold 10^−15^ was treated as extinct and removed from the dynamical system. The numbers of surviving bacterial phenotypes and phage species at the final integration time were denoted by *N*_*s*_ and *M*_*s*_, respectively.

For each parameter set, we generated independent random trait realizations and computed the corresponding long-time survivor counts. The phage diversity reported in the main text was quantified by the ensemble-averaged number of surviving phage species,

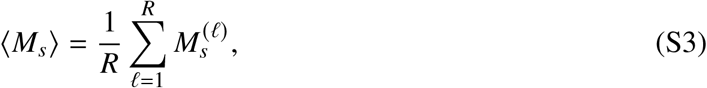

where *R* is the number of independent realizations and 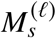 is the number of surviving phage species in realization ℓ. For the experimentally motivated two-phage setting in Fig. 1(e) of the main text, ⟨*M*_*s*_⟩_{1+*η*, 1, 1−*η*}_ was additionally averaged over the three host-growth regimes with mean growth rates ⟨*r*⟩ = 1 + *η*, 1, and 1 − *η*, as described in the corresponding figure caption.

Because convergence times varied across random trait realizations and parameter regimes, we calibrated the final integration time using long-time reference simulations. Specifically, for representative parameter sets, we integrated the dynamics up to a reference time of *T* = 10^8^ and recorded the numbers of surviving bacterial phenotypes and phage species at intermediate checkpoints. We then compared these intermediate survivor counts with the corresponding values at *T* = 10^8^. The final integration times were chosen so that both *N*_*s*_ and *M*_*s*_ agreed with the *T* = 10^8^ reference values in at least 94% of realizations.

Convergence was slower in regimes with relatively low trait heterogeneity. We therefore used different final integration times for low- and high-heterogeneity regimes, with the precise cutoffs and integration times determined separately for each model class and trait distribution. For the experimentally motivated two-phage setting shown in Fig. 1 of the main text, we used final integration time *T* = 10^7^ for *σ*_*r*_ < 0.05 and *T* = 2 × 10^6^ for *σ*_*r*_ ≥ 0.05. The detailed cutoffs, final integration times, and ensemble sizes used for other analysis are given in Supplementary Text.

#### Infection kernels and trait distributions

In the main text, we used a Gaussian infection kernel and Gaussian trait distributions. For the Gaussian infection kernel, we denote the kernel width by 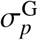, which corresponds to *σ*_*p*_ in the main text. For Gaussian trait distributions, we denote the trait dispersion by *σ*^G^, which corresponds to *σ* in the main text.

Throughout this section, *r*_*i*_ denotes the growth rate of bacterial phenotype *i*, and *q*_*α*_ denotes the preferred host growth state of phage species *α*. We write *A*(*r, q*_*α*_) for the continuous infection kernel of phage *α* as a function of host growth rate *r*. The affinity matrix used in the dynamical equations is obtained by evaluating this infection kernel at the sampled bacterial growth rates,

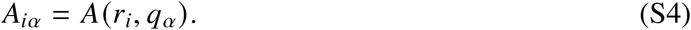

##### Infection kernels

For the Gaussian infection kernel, we used

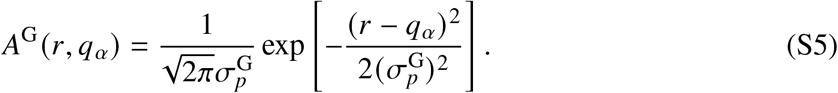

Here, 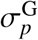 controls the phage niche width in growth-rate space. The infection efficiency is maximal at *r* = *q*_*α*_ and decays symmetrically as the mismatch between the bacterial growth rate and the phage preferred host state increases. This is the infection kernel used in the main text, where we write 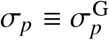.

##### Trait distributions

In the generalized bacteria–phage ensembles, bacterial growth rates *r*_*i*_ and phage preferred host states *q*_*α*_ were sampled independently from the same type of trait distribution. Thus, for a given trait distribution *p* (*x*), we sampled

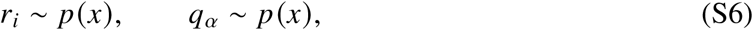

independently for all *i* = 1, …, *N* and *α* = 1, …, *M*. This matched host–phage trait ensemble allows us to vary the common scale of host phenotypic dispersion and phage preference dispersion while keeping the characteristic trait scale fixed around 1.

For Gaussian trait distributions, we used a normal distribution with parent mean *m* and parent standard deviation *σ*^G^, restricted to positive trait values:

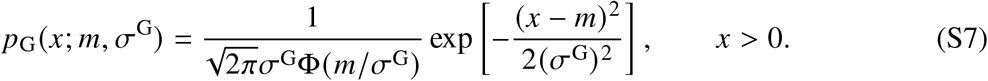

Here, Φ is the cumulative distribution function of the standard normal distribution. This distribution is equivalent to sampling from *N* (*m*, (*σ*^G^)^2^) and redrawing whenever a negative value is obtained. The parameter *σ*^G^ therefore controls the width of the parent Gaussian distribution. In the generalized ensembles of the main text, we used *m* = 1 and write *σ* ≡ *σ*^G^. In the experimentally motivated two-phage setting, bacterial growth rates were sampled from the same positive-truncated Gaussian distribution with parent means *m* = 1 + *η*, 1, and 1 − *η*, corresponding to the high-, intermediate-, and low-growth host populations, respectively, while the two phage preferred host states were fixed at *q*_1_ = 1 + *η* and *q*_2_ = 1 − *η*.

#### Niche differentiation (ND) and invasion viability (IV)

To quantify the mechanism underlying the optimal coexistence window, we introduced two quantities: niche differentiation (ND) and invasion viability (IV). ND measures the expected separation between phage infection kernels along the positive host growth-rate axis, whereas IV measures the probability that a randomly drawn host–phage pair satisfies a pairwise rare-invasion condition. These quantities are not intended to provide necessary or sufficient conditions for coexistence in the full multispecies community. Rather, they serve as mechanistic indicators of the two opposing constraints that shape phage coexistence: niche separation among phages and the availability of viable host–phage infection links.

##### Niche differentiation

Let *A*(*r, q*_*α*_) denote the continuous infection kernel of a phage species *α* whose preferred host growth state is *q*_*α*_. The affinity matrix used in the dynamical equations is obtained by evaluating this kernel at the sampled bacterial growth rates, *A*_*iα*_ = *A*(*r*_*i*_, *q*_*α*_), as defined in Eq. (S4).

For two phages with preferred host growth states *q*_*α*_ and *q*_*β*_, we define their pairwise niche differentiation as the normalized non-overlapping area between their infection kernels,

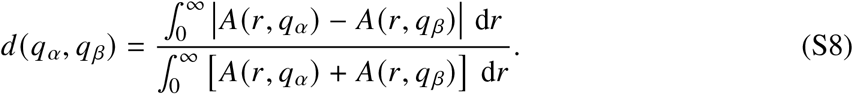

This definition measures kernel separation on the biologically relevant domain of positive host growth rates. By construction, 0 ≤ *d* (*q*_*α*_, *q*_*β*_) ≤ 1. The value is zero when the two kernels are identical and approaches one when their overlap is negligible.

The expected niche differentiation at a given trait parameter and kernel parameter is defined by averaging this pairwise separation over two independently sampled phage preferred host states:

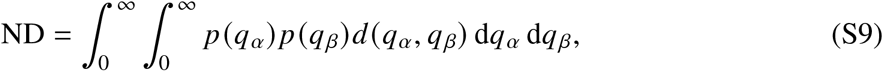

where *p* (*q*) is the chosen trait distribution defined above. Thus, ND is an expected pairwise niche separation under the phage preference distribution.

For the Gaussian infection kernel, Eq. (S8) can be evaluated analytically on the positive growth-rate domain. Because the pairwise niche differentiation is symmetric under exchange of the two phages, we define

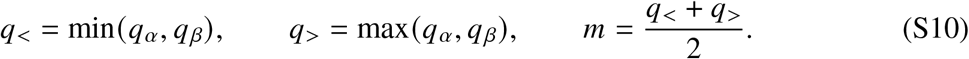

If *q*_*α*_ = *q*_*β*_, then *d*_G_(*q*_*α*_, *q*_*β*_) = 0. Otherwise, we define

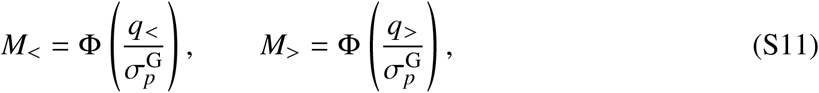

and

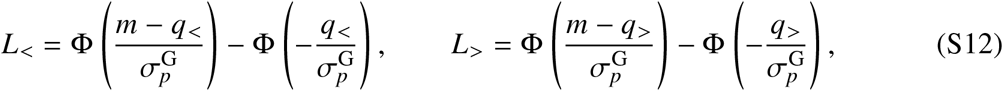

where Φ is the cumulative distribution function of the standard normal distribution. The pairwise niche differentiation for the Gaussian infection kernel is then

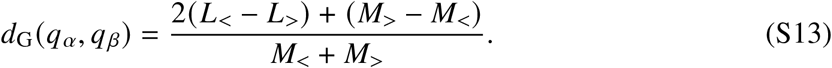

This expression is the normalized non-overlapping area of two Gaussian infection kernels restricted to *r* > 0.

##### Invasion viability

IV quantifies the probability that a randomly drawn host–phage pair satisfies a pairwise rare-invasion condition. Consider a single bacterial phenotype with growth rate *r* in the absence of phages. From Eq. (S1), its positive resource-limited equilibrium is

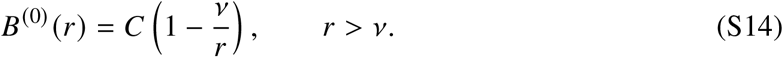

If a phage with preferred host growth state *q* is introduced at low density, its initial per-capita growth rate is

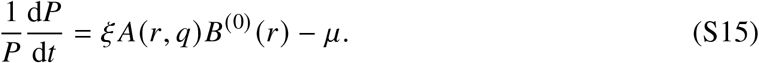

Therefore, the phage can invade if

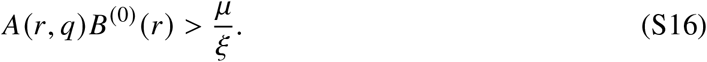

The following expressions are written for the numerical choice *C* = 1.

We define invasion viability as the probability that Eq. (S16) is satisfied when the host growth rate *r* and the phage preferred host state *q* are independently sampled from the chosen trait distribution:

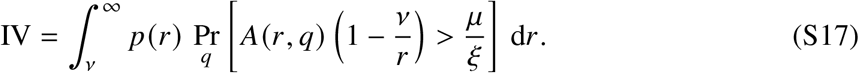

Equivalently,

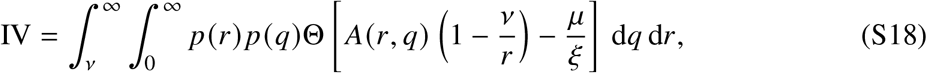

where Θ(*x*) is the Heaviside step function. Thus, IV ranges from 0 to 1 and represents the expected fraction of viable pairwise host–phage links under the trait distribution.

For the Gaussian infection kernel,

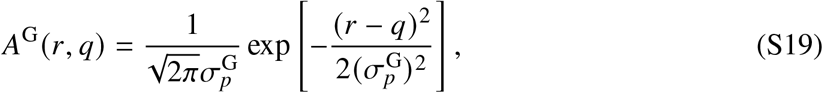

the invasion condition is satisfied only if

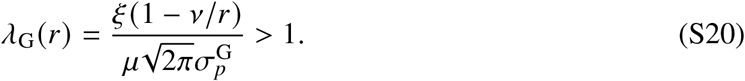

When this condition holds, viable phage preferences satisfy

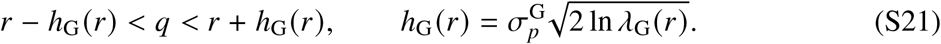

Therefore,

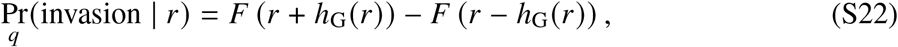

where *F* is the cumulative distribution function of the chosen phage preference distribution. If *λ*_G_(*r*) ≤ 1, this probability is set to zero.

For the positive-truncated Gaussian trait distribution used in the main text, with parent mean 1 and standard deviation *σ*^G^, the cumulative distribution function is

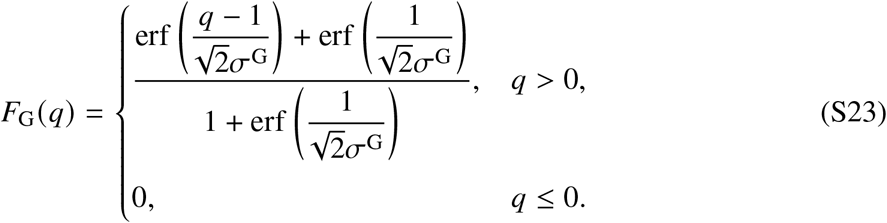

Defining

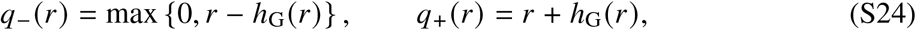

the conditional invasion probability for *λ*_G_(*r*) > 1 becomes

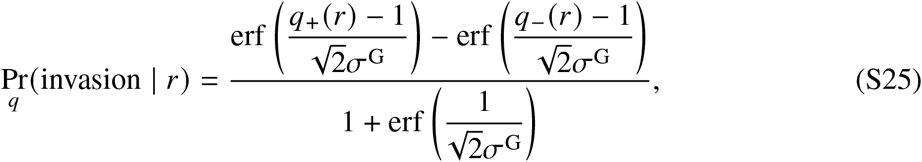

and it is zero for *λ*_G_(*r*) ≤ 1. Thus, for the Gaussian trait distribution used in the main text,

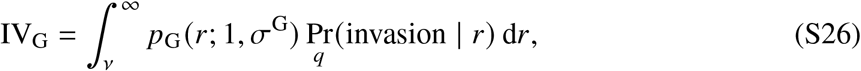

where the conditional invasion probability is given by Eq. (S25) when *λ*_G_(*r*) > 1 and is zero otherwise.

## Supplementary Text

### Robustness across model specifications

In the main text, we focused on communities with *N* = *M* = 10, Gaussian infection kernels, Gaussian trait distributions, and demographic parameters *ξ* = 2, *μ* = 1, and *v* = 0.1. Here, we test whether the optimal coexistence window and its interpretation in terms of niche differentiation (ND) and invasion viability (IV) persist when we vary (i) the infection kernel and trait distributions, (ii) the demographic parameters, (iii) the initial numbers of bacterial phenotypes and phage species, and (iv) the phenotypic inheritance rule during bacterial reproduction.

#### Robustness across infection kernels and trait distributions

We first tested whether the optimal coexistence window depends on the functional forms used to describe bacterial and phage traits or infection kernels. For generalized communities with *N* = *M* = 10, we considered all combinations of Gaussian and log-normal infection kernels with Gaussian, log-normal, and uniform trait distributions. The Gaussian infection kernel and Gaussian trait distribution are defined in the Materials and Methods section of the Supplementary Information. For the additional model specifications considered here, we denote the log-space width of the log-normal infection kernel by 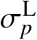, the log-space dispersion of the log-normal trait distribution by *σ*^L^, and the half-width of the uniform trait distribution by *k*.

For the log-normal infection kernel, we used

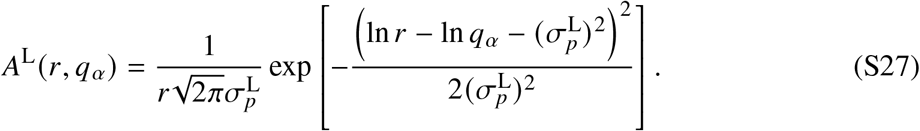

The parameter 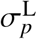 controls the width of the infection kernel in log-growth-rate space. The shift 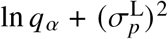 is chosen so that the mode of the log-normal kernel occurs at *r* = *q*_*α*_. Thus, as in the Gaussian case, the infection efficiency is maximal when *r* = *q*_*α*_, and *q*_*α*_ represents the phage’s preferred host growth state.

For log-normal trait distributions, we used

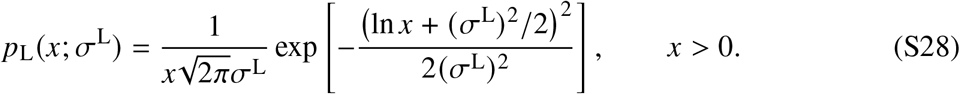

Equivalently, ln *x* is normally distributed with mean −(*σ*^L^)^2^/2 and variance (*σ*^L^)^2^. With this parameterization, the mean trait value is

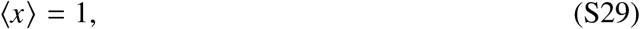

and the variance is

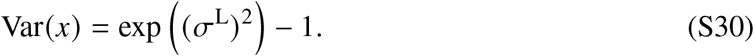

Thus, *σ*^L^ controls the dispersion in log-trait space while keeping the mean trait value fixed at 1.

For uniform trait distributions, we used

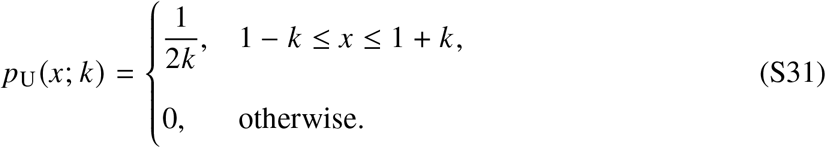

Here, *k* is the half-width of the uniform distribution. The mean and variance are

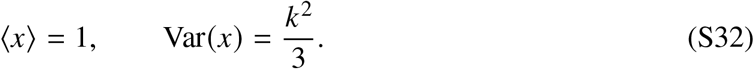

All values of *k* used in this study satisfy 0 < *k* < 1, ensuring that all sampled trait values remain positive.

For Gaussian and log-normal trait distributions, the final integration time was set to *T* = 10^7^ when the corresponding trait-dispersion parameter was smaller than 0.1 and to *T* = 2 × 10^6^ otherwise. For uniform trait distributions, we used *T* = 10^7^ for *k* < 0.25 and *T* = 2 × 10^6^ for *k* ≥ 0.25.

Across all six combinations, ⟨*M*_*s*_⟩ remains low when the trait dispersion is too small relative to the infection-kernel width and also decreases when the trait dispersion becomes too large relative to a narrow infection kernel [Fig. S1]. The precise location and magnitude of the coexistence maximum depend on the kernel and trait distribution, but a region of enhanced phage richness consistently appears at intermediate scale matching. The distributions of *M*_*s*_ at representative parameter sets also retain the qualitative distinction observed in the main text. Small-dispersion, broad-kernel regimes generally produce relatively narrow distributions with low but typically nonzero phage richness, whereas large-dispersion, narrow-kernel regimes produce broader distributions containing more realizations with few or no surviving phages.

We next examined whether the ND–IV interpretation remains valid across these model specifications. For the log-normal infection kernel, the kernels are supported on *r* > 0 and normalized on the positive domain. The pairwise niche differentiation reduces to

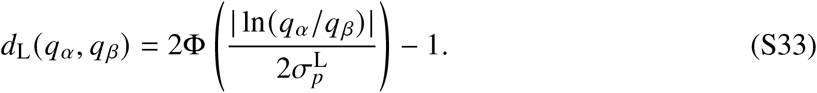

For the log-normal infection kernel,

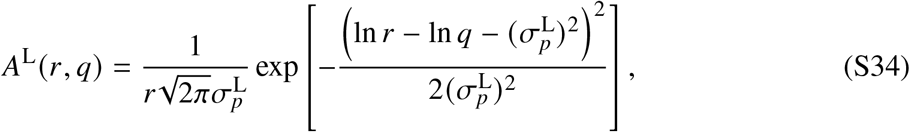

the invasion condition is satisfied only if

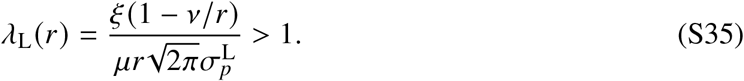

When this condition holds, viable phage preferences satisfy

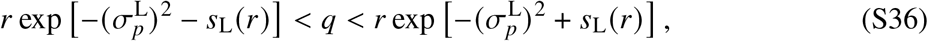

where

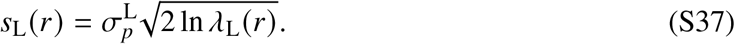

Therefore,

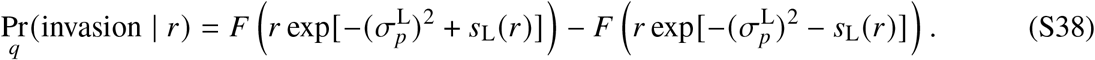

If *λ*_L_(*r*) ≤ 1, this probability is set to zero. Here, *F* denotes the cumulative distribution function of the corresponding phage preference distribution.

For the Gaussian infection kernel, ND increases as the trait dispersion becomes larger relative to the infection-kernel width, whereas IV decreases because an increasing fraction of host–phage pairs fails to provide sufficient infection-weighted host biomass for invasion [Fig. S2]. The maximum of ⟨*M*_*s*_⟩ lies between these two limiting regimes for Gaussian, log-normal, and uniform trait distributions.

The same opposing trends are observed for the log-normal infection kernel [Fig. S3]. Despite the asymmetry of this kernel, ND increases with the relative trait dispersion while IV decreases, and phage richness is again maximized over an intermediate range of the corresponding scale ratio. The optimal coexistence window is therefore not specific to either the symmetry of the Gaussian kernel or the shape of the trait distribution.

#### Robustness across demographic parameters

We next varied the demographic parameters governing phage production, phage loss, and bacterial loss. These analyses used *N* = *M* = 10, Gaussian trait distributions, and the Gaussian infection kernel. For all demographic-parameter scans, the final integration time was set to *T* = 10^7^ when *σ*^G^ < 0.1 and to *T* = 2 × 10^6^ otherwise.

We first varied the phage burst-size parameter among *ξ* = 2, 5, and 10, while fixing *μ* = 1 and *v* = 0.1. Changing *ξ* produces only minor quantitative differences in the phage-richness landscape, and the intermediate coexistence window remains qualitatively similar across the tested values [Fig. S4]. ND is identical for all values of *ξ* because it depends only on the trait distributions and infection-kernel geometry, whereas IV increases only slightly with *ξ* owing to the relaxed pairwise invasion threshold [Fig. S5]. In all cases, ⟨*M*_*s*_⟩ reaches its highest values in the intermediate region where ND is sufficiently high while IV remains sufficiently large, supporting the same trade-off mechanism identified in the main text.

We then varied the phage degradation rate among *μ* = 0.2, 0.5, and 1, while fixing *ξ* = 2 and *v* = 0.1. Changing *μ* likewise produces only minor quantitative differences in the phage-richness landscape, and the intermediate coexistence window remains qualitatively similar across the tested values [Fig. S6]. ND is identical for all values of *μ*, whereas IV decreases modestly with increasing *μ* owing to the more restrictive pairwise invasion threshold [Fig. S7]. For all tested degradation rates, ⟨*M*_*s*_⟩ is maximized where sufficient niche differentiation coexists with sufficiently high invasion viability.

Finally, we varied the bacterial dilution or mortality rate among *v* = 0.02, 0.05, and 0.1, while fixing *ξ* = 2 and *μ* = 1. Changing *v* again produces only minor quantitative differences in the phage-richness landscape, while the intermediate coexistence window remains qualitatively similar across the tested values [Fig. S8]. ND is independent of *v*, whereas IV decreases modestly with increasing *v* because the phage-free bacterial biomass available to support invasion is reduced [Fig. S9]. For all tested bacterial loss rates, ⟨*M*_*s*_⟩ is maximized in the intermediate region where niche differentiation is sufficiently strong while invasion viability remains sufficiently high.

Together, these parameter scans show that the nonmonotonic coexistence window is robust to variation in the demographic rates over the ranges examined. Although *ξ, μ*, and *v* produce modest changes in IV through the pairwise phage-invasion condition, they do not alter ND or the central result that enhanced phage richness occurs where niche differentiation and invasion viability are jointly maintained.

#### Robustness across community sizes

We next tested whether the coexistence window persists when the initial numbers of bacterial phenotypes and phage species are varied. Using Gaussian infection kernels and Gaussian trait distributions, we considered *N* = *M* = 5, *N* = *M* = 10, and *N* = *M* = 20. For *N* = *M* = 5 and *N* = *M* = 10, the final integration time was set to *T* = 10^7^ for *σ*^G^ < 0.1 and to *T* = 2 × 10^6^ for *σ*^G^ ≥ 0.1. For *N* = *M* = 20, we used *T* = 2 × 10^7^ for all parameter sets.

The absolute number of surviving phages increases with the number of species initially available, as expected, and the distribution of *M*_*s*_ changes quantitatively with community size. Nevertheless, all three community sizes display the same qualitative organization of parameter space: phage richness is suppressed when trait dispersion is either too small or too large relative to the infection-kernel width and is enhanced over an intermediate range [Fig. S10]. Thus, the optimal coexistence window is not a finite-size feature of the *N* = *M* = 10 ensemble used in the main text.

#### Robustness to the assumption of phenotypic inheritance

In the main model, each bacterial phenotypic state is treated as a persistent subpopulation over the ecological timescale of the dynamics. This represents one limiting case of phenotypic inheritance. At the opposite extreme, phenotypic memory may be completely lost upon reproduction, such that the phenotype of a newly produced bacterial cell is independent of that of its parent.

To test whether our main result depends on the assumption of persistent phenotypic states, we considered an alternative model in which each newly produced bacterial cell is assigned uniformly at random to one of the *N* predefined phenotypic states. Under this assumption, the bacterial dynamics become

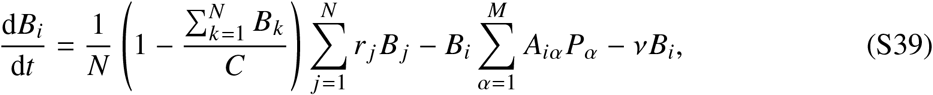

whereas the phage dynamics remain unchanged:

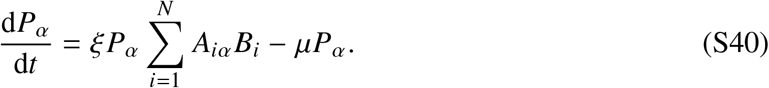

Thus, the total bacterial production rate is unchanged relative to the main model, but newly produced biomass is redistributed uniformly among the predefined phenotypic states.

Because bacterial phenotypic states can be continuously regenerated in this model, a bacterial phenotype that reaches a very low abundance is not permanently removed from the dynamics.

Phage species, by contrast, remain absorbing upon extinction and were treated as extinct when their density fell below the same threshold used in the main model.

For this alternative model, we calibrated the final integration time using a reference time of *T* = 10^7^. For representative parameter sets, we recorded the number of surviving phage species *M*_*s*_ at intermediate checkpoints and compared these values with the corresponding survivor counts at *T* = 10^7^. The final integration times were chosen such that *M*_*s*_ agreed with the *T* = 10^7^ reference value in at least 95% of realizations. We used *T* = 5 × 10^5^ for *σ* < 0.1 and *T* = 10^5^ for *σ* ≥ 0.1.

Despite this substantial change in the phenotypic inheritance rule, the qualitative coexistence pattern remains unchanged [Fig. S11]. Phage richness remains low when trait dispersion is small relative to the infection-kernel width, increases at intermediate scale matching, and decreases again when the trait dispersion becomes too large relative to a narrow infection kernel. The distribution of *M*_*s*_ likewise retains the qualitative distinction between the two low-diversity regimes. Thus, the existence of an intermediate coexistence optimum does not rely on perfect persistence of bacterial phenotypic states across reproduction.

Taken together, these analyses show that the nonmonotonic relationship between phenotypic dispersion and phage richness is robust within the range of model structures and parameters examined here. Changing the infection kernel, trait distribution, demographic rates, community size, or phenotypic inheritance rule alters quantitative details but preserves the central requirement that phage niches be sufficiently differentiated while viable host–phage links remain sufficiently abundant.

### Minimal two-host–two-phage model

Finally, we consider a minimal community composed of two bacterial phenotypes and two phage species. Unlike the model with general bacteria-phage ensembles, we can solve this model analytically. Inspired by the trait-based structure of the generalized bacteria–phage ensembles, we construct a structured 2 × 2 affinity matrix in which each phage preferentially infects one of the two host states. We derive the feasible and invasion-resistant equilibrium phases, their phase boundaries, and the corresponding average phage richness, niche differentiation, and invasion viability.

#### Structured two-host–two-phage model

In the minimal model, the state of the system at time *t* is described by the vector 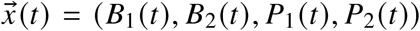, where the first (last) two components refer to the bacteria (phage) abundances. For bacterial phenotype *i* (*i* ∈ {1, 2}), the time evolution of the abundance reads

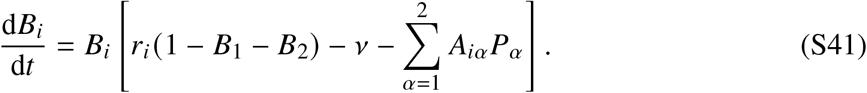

Analogously, the abundance of phage *α* (*α* ∈ {1, 2}) is governed by:

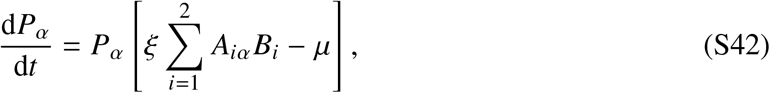

where the carrying capacity *C* has been set to 1.

The two bacterial growth states are positioned symmetrically around 1:

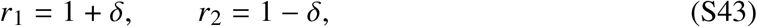

where 2*δ* is their separation. The preferred host states of the two phages are

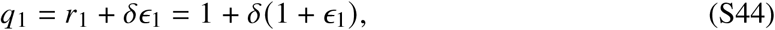

and

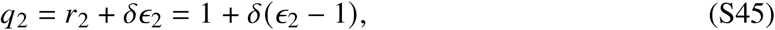

with

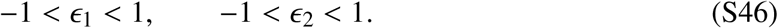

Consequently,

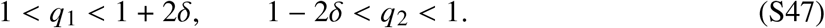

Thus, *P*_1_ is constructed to preferentially infect the faster-growing state *B*_1_, whereas *P*_2_ preferentially infects the slower-growing state *B*_2_.

Using the Gaussian infection kernel, we define

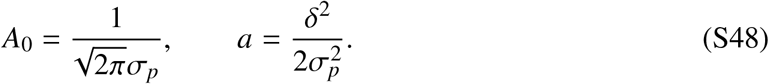

The four affinity-matrix elements are

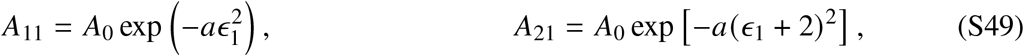

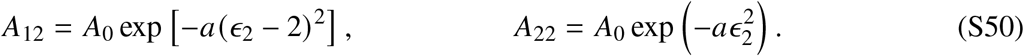

For −1 < *ϵ*_1_, *ϵ*_2_ < 1, the host–phage trait mismatches satisfy

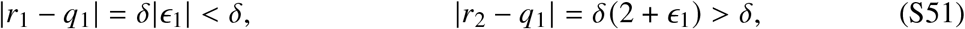

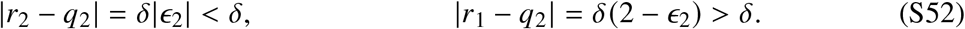

The former choices thus guarantee that bacteria 1 (2) represents the ecological niche of phage 1 (2). Mathematically:

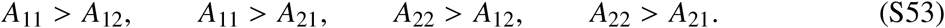

For convenience, we define

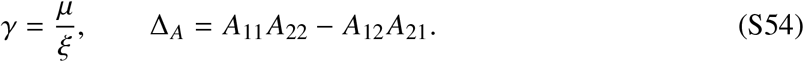

The affinity ordering in Eq. (S53) implies

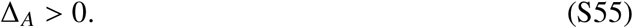

#### Equilibrium phases and exclusion of alternative phases

First, we derive the different steady states of the system. Assuming that both bacteria phenotypes and phages species are in the system, the fixed points of the dynamics are:

- Bacteria-only phase: 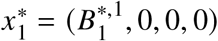,
- Bacteria and its phage: 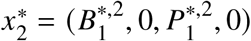,
- Phage killing the winner: 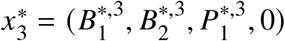,
- Full coexistence: 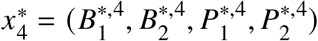,

where the superscript denotes the associated fixed point for each variable. The other fixed points of the Eqs. (S41) and (S42) are stable only when communities misses one bacteria phenotype or phage species. Specifically, the fixed point 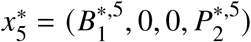 is unstable to invasion by phage 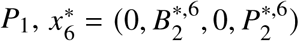 is unstable to invasion by bacterial phenotype *B*_1_, and 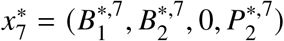 is unstable to invasion by phage *P*_1_.

There is a single stable fixed point for each combination of the parameters of the model (*σ*_*p*_, *δ, ϵ*_1_, *ϵ*_2_). In Fig. S12 we show multiple phase diagrams corresponding to different choices of *σ*_*p*_ and *δ*. In what follows, we focus on determining analytically the boundaries between the different fixed points (phases) of our system. For brevity, in each sub-subsection analyzing the stability of a given fixed point *m*, we skip the superscript associating with the fixed point, i.e. we assume 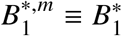.

#### Equilibrium conditions and phase boundaries

*Bacteria-only phase:* 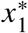: Because *r*_1_ > *r*_2_, the bacteria-only equilibrium contains the faster-growing state:

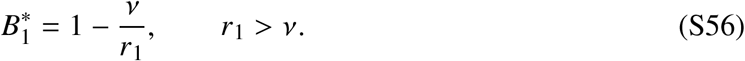

Note that if *v* > *r*_1_, the equilibrium of the system will be the extinction of both bacteria and phages. Restricting to *v* < *r*_1_, the invasion growth rate of *B*_2_ is

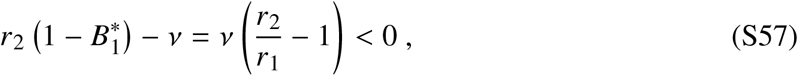

so coexistence between both bacteria phenotypes without phages is forbidden. Now we compute the stability criteria for 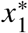 with respect to phages. Because *A*_11_ > *A*_12_, the *B*_1_ phase is resistant to both phages when

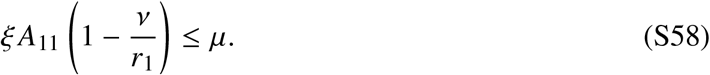

The boundary between 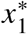 and 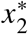 is therefore

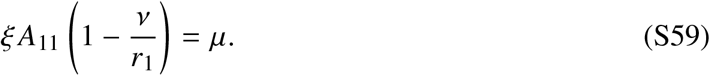

Defining

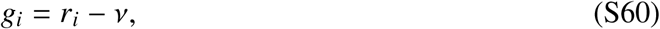

the two possible solutions are

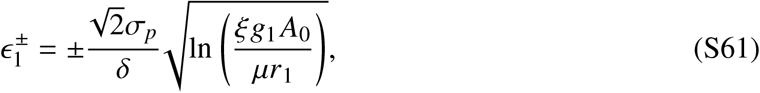

provided that the logarithm is positive and the roots lie within (−1, 1).

##### Bacteria and its phage: 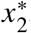

The equilibrium densities are

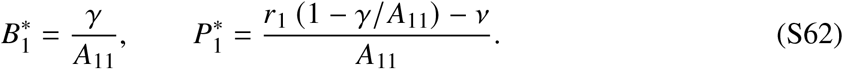

This equilibrium is feasible when

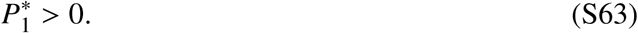

The absent phage *P*_2_ cannot invade because

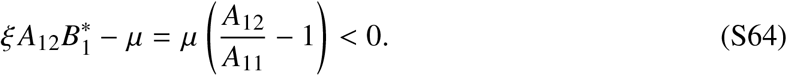

The transition from 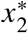 to 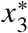 occurs when the invasion growth rate of *B*_2_ becomes zero:

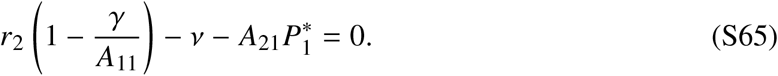

Substituting Eq. (S62) gives

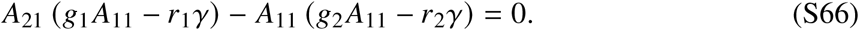

Because *A*_11_ and *A*_21_ depend only on *ϵ*_1_, this boundary is independent of *ϵ*_2_ and appears as a vertical line in the (*ϵ*_1_, *ϵ*_2_) plane. Only roots for which the resident *B*_1_*P*_1_ equilibrium is feasible correspond to physical phase boundaries.

##### Phage killing the winner 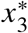

Writing

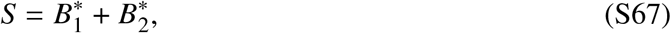

the bacterial equilibrium conditions yield

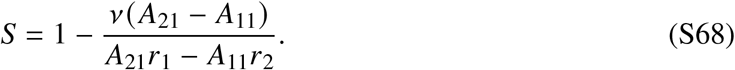

The individual equilibrium densities are

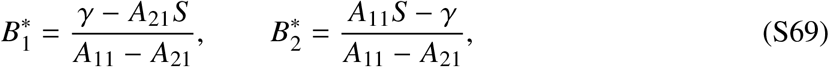

and the phage 1 abundance reads:

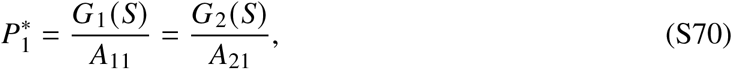

where we have defined *G*_*i*_ (*S*) = *r*_*i*_ (1 − *S*) − *v*. This phase requires

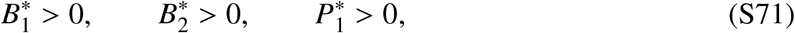

and resistance to invasion by *P*_2_:

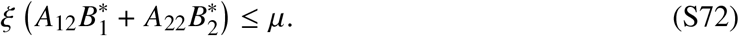

Therefore, the boundary between 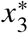 and 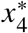 is determined by equality in Eq. (S72):

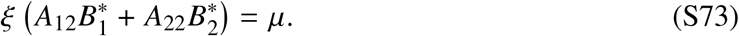

Defining

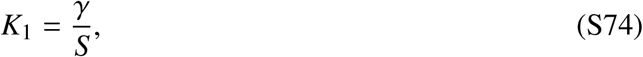

this condition can be written as

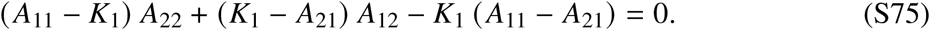

Because *A*_11_ and *A*_21_ depend on *ϵ*_1_, whereas *A*_12_ and *A*_22_ depend on *ϵ*_2_, this equation defines the curved boundary in the (*ϵ*_1_, *ϵ*_2_) plane.

##### Full coexistence: 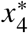

The two phage equilibrium conditions give

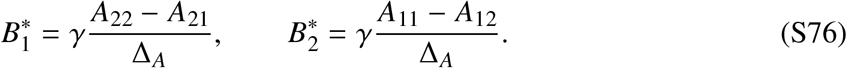

Both bacterial densities are positive because of Eq. (S53) and Δ_*A*_ > 0. Writing 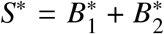, the phage densities are

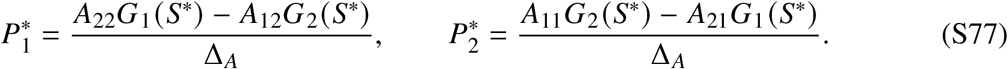

The full-coexistence phase is feasible when

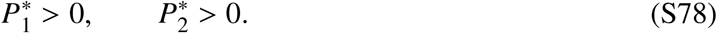

#### Average phage richness

To reproduce the figures shown in the manuscript in this minimal model, we fix *σ*_*p*_ and *δ* and compute the average phage diversity ⟨*M*_*s*_⟩ observed across the entire phase diagram delimited by *ϵ*_1_ and *ϵ*_2_, as sketched in Fig. S13(a). Treating *ϵ*_1_ and *ϵ*_2_ as independent random variables uniformly distributed on (−1, 1), the average number of surviving phages is

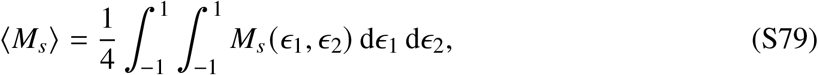

where *M*_*s*_ = 0, 1, or 2 depending on the equilibrium phase. If *f*_*m*_ is the fraction of the phase diagram occupied by states with *m* surviving phages, then

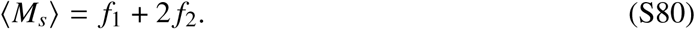

Fig. S13(b) shows that phage diversity depends non-monotonically on the phenotypic heterogeneity *δ* and the phage specialization *σ*_*p*_. Therefore, the two-bacteria-two-phage minimal model allows us to retrieve analytically the phenomenology reported in the main manuscript for more complex bacteria-phage ensembles. Next, we compute the niche differentiation and the invasion viability analytically to check whether the optimal diversity is related to the ecological trade-off described in the main text.

#### Niche differentiation and invasion viability

Because *q*_1_ > *q*_2_ throughout the parameter domain, the pairwise niche differentiation between the two Gaussian infection kernels is

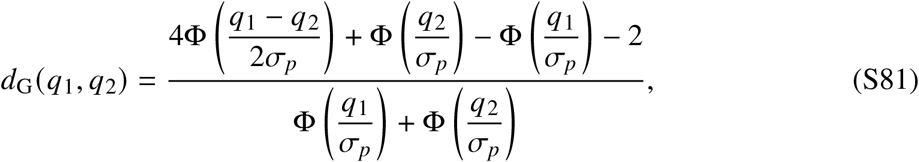

where Φ is the standard-normal cumulative distribution function. The niche differentiation is

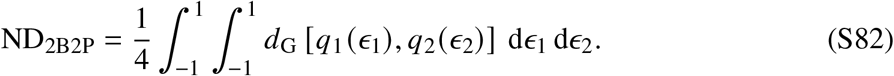

Because the model is explicitly constructed such that *P*_1_ specializes on *B*_1_ and *P*_2_ specializes on *B*_2_, we can compute invasion viability just considering the pairs

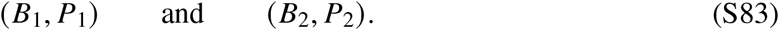

Cross-infection remains fully included in the population dynamics but is excluded from this specialized IV average.

For host state *i*, the corresponding phage-free single-host density is

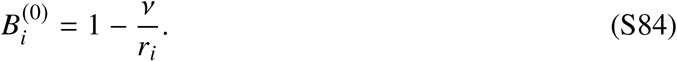

The matched pair (*B*_*i*_, *P*_*i*_) is viable when

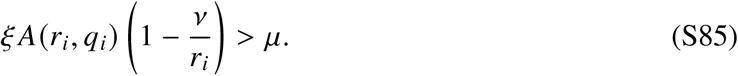

Defining

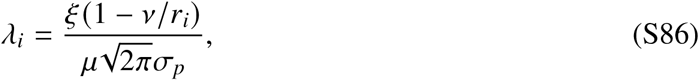

the viable interval exists when *λ*_*i*_ > 1 and is given by

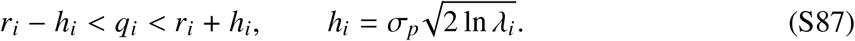

Let *F*_*i*_ (*q*) denote the cumulative distribution function of

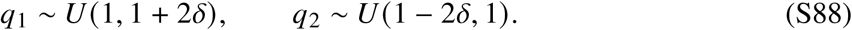

The viability probability of the matched pair is

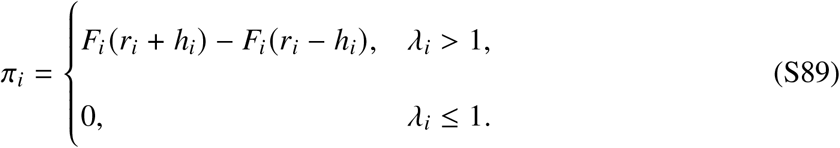

The specialized invasion viability is therefore

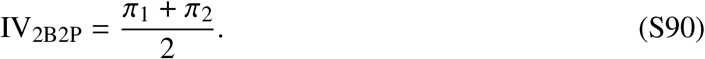

This pairwise diagnostic is evaluated in the corresponding single-host phage-free equilibrium and is not itself a stability condition for the full four-population community.

Fig. S13(c)–(e) reproduces analytically the ecological trade-off caused by promoting phenotypic heterogeneity as ND increases (Fig. S13(c)), hampering the potential competition among phages, but IV decreases (Fig. S13(d)) as a result of the mismatch between the growth rate of the bacteria and the niches of both phages. Full phage coexistence is therefore most likely when niche separation is sufficiently strong while matched-pair invasion viability remains sufficiently high (Fig. S13(e)).

**Figure S1:**
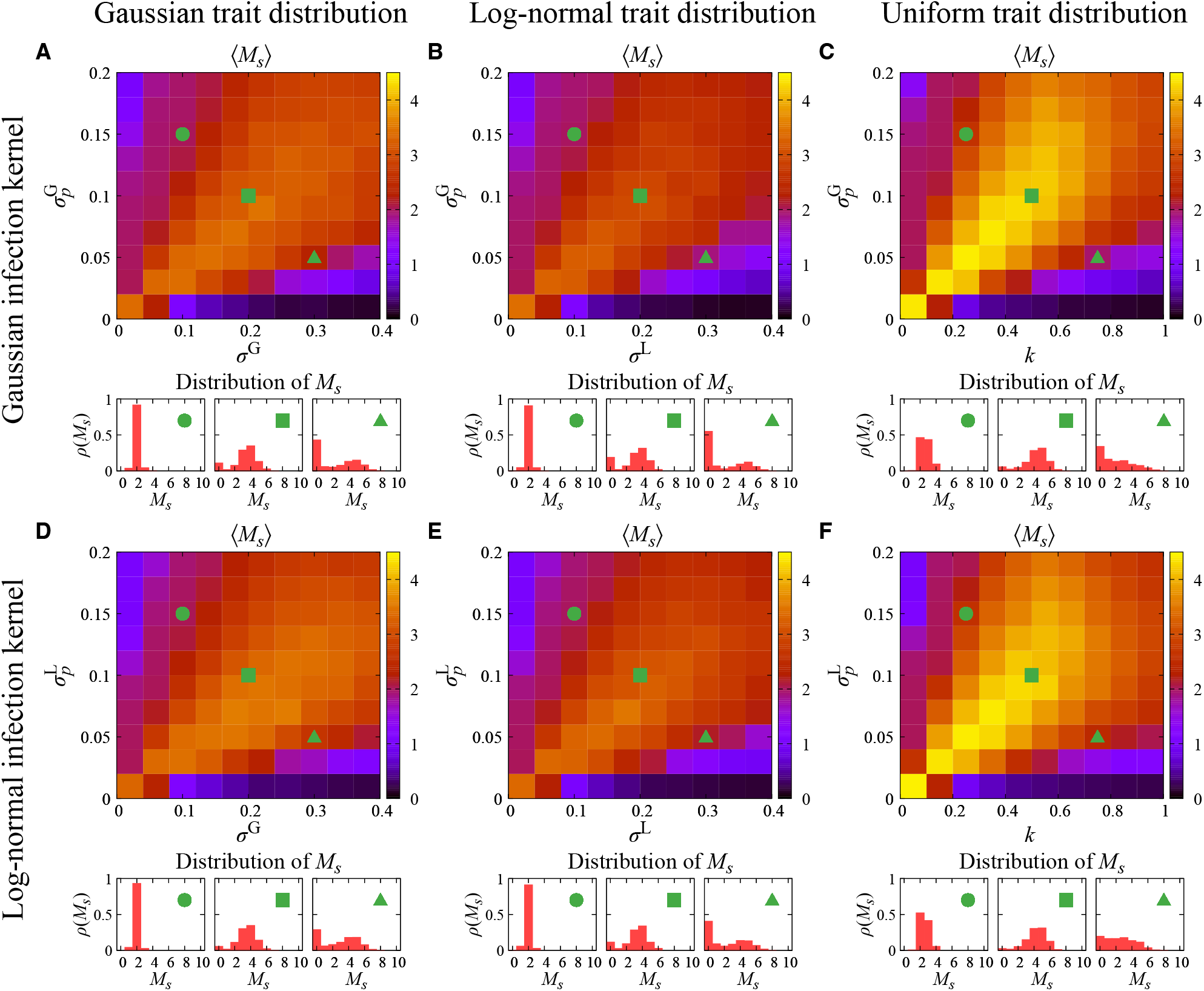
Robustness of phage diversity across infection kernels and trait distributions. Average number of surviving phage species ⟨*M*_*s*_⟩ and distributions of *M*_*s*_ for generalized bacteria– phage ensembles with *N* = *M* = 10. The top and bottom rows correspond to the Gaussian and log-normal infection kernels, respectively. Columns correspond to Gaussian trait distributions (**A, D**), log-normal trait distributions (**B, E**), and uniform trait distributions (**C, F**). The horizontal axes show the corresponding trait-dispersion parameters: *σ*^G^ for Gaussian traits, *σ*^L^ for log-normal traits, and *k* for uniform traits. The vertical axes show the infection-kernel widths 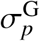 and 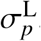. The lower histograms show the distributions of *M*_*s*_ at the three representative parameter sets indicated by the green symbols. Results are averaged over 10^3^ independent trait realizations.

**Figure S2:**
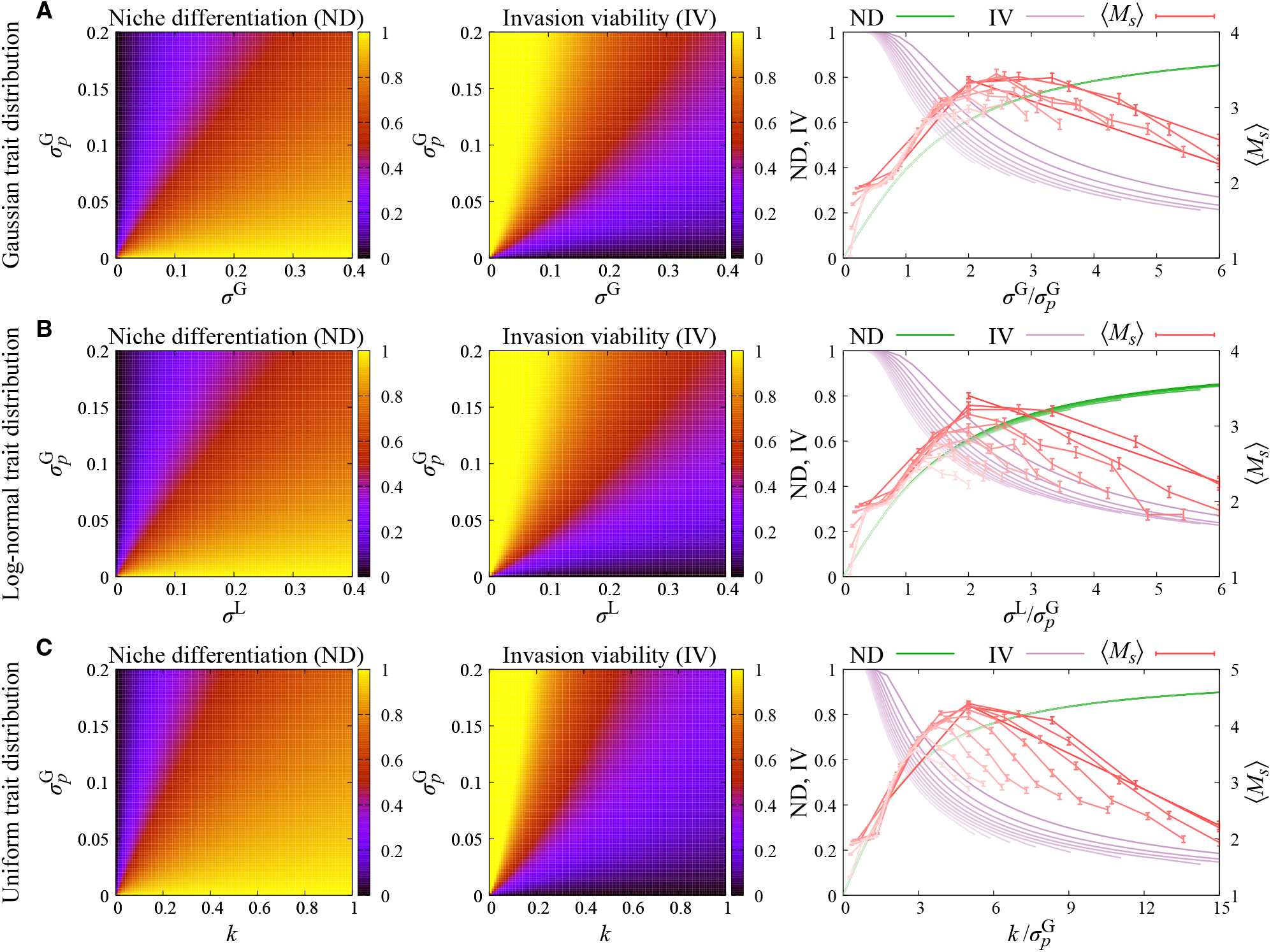
Niche differentiation and invasion viability for the Gaussian infection kernel across trait distributions. ND, IV, and ⟨*M*_*s*_⟩ for the Gaussian infection kernel combined with (**A**) Gaussian, (**B**) log-normal, and (**C**) uniform trait distributions. In each row, the left and middle panels show ND and IV, respectively, and the right panel shows ND, IV, and ⟨*M*_*s*_⟩ as functions of the corresponding scale ratio. The scale ratios are 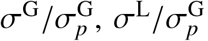, and 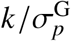. Each curve corresponds to a fixed kernel width 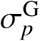 from 0.01 to 0.19 in increments of 0.02. Error bars for ⟨*M*_*s*_⟩ denote standard errors across independent trait realizations.

**Figure S3:**
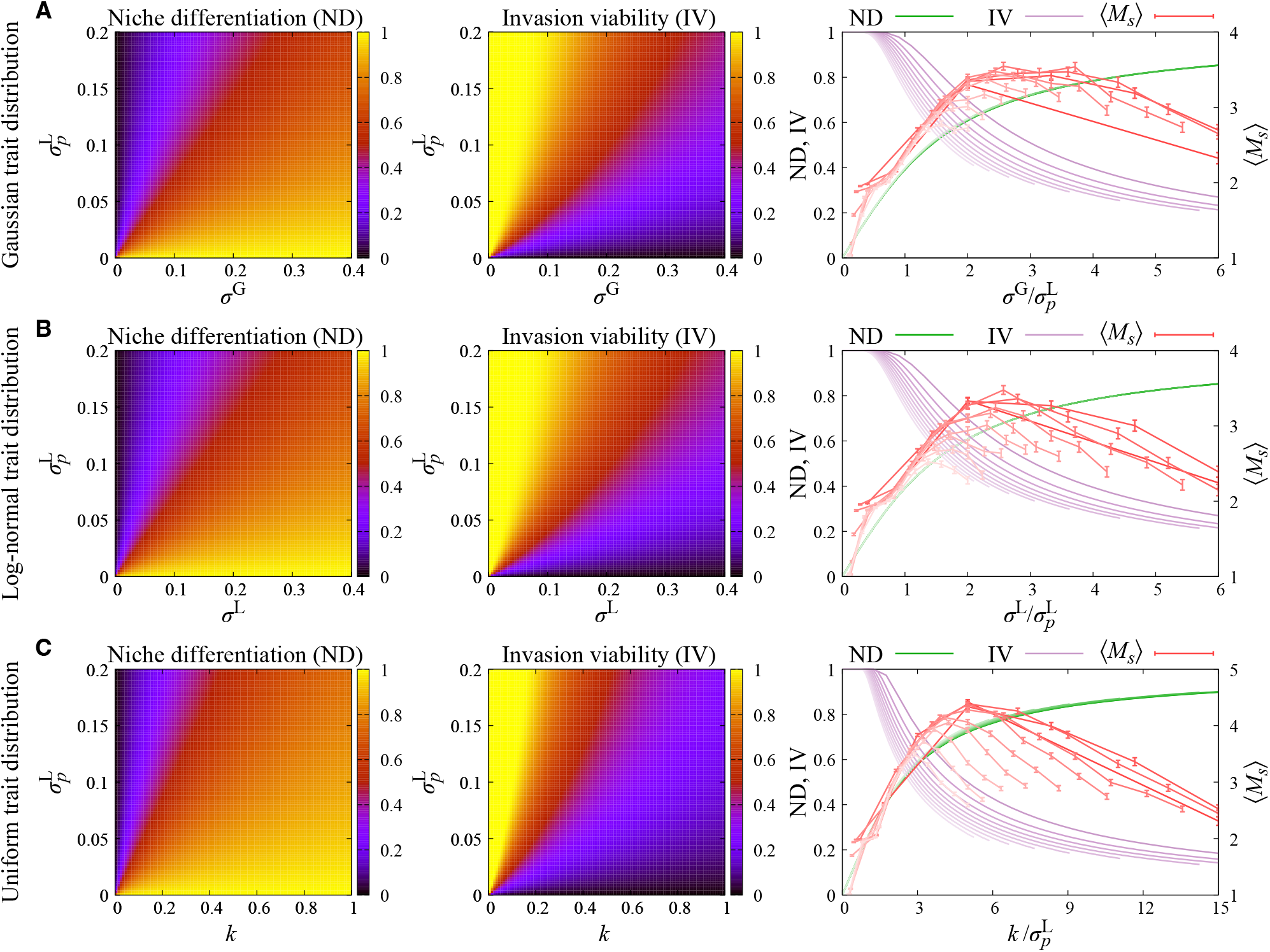
Niche differentiation and invasion viability for the log-normal infection kernel across trait distributions. ND, IV, and ⟨*M*_*s*_⟩ for the log-normal infection kernel combined with (**A**) Gaussian, (**B**) log-normal, and (**C**) uniform trait distributions. In each row, the left and middle panels show ND and IV, respectively, and the right panel shows ND, IV, and ⟨*M*_*s*_⟩ as functions of the corresponding scale ratio. The scale ratios are 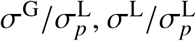, and 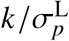. Each curve corresponds to a fixed kernel width 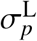 from 0.01 to 0.19 in increments of 0.02. Error bars for ⟨*M*_*s*_⟩ denote standard errors across independent trait realizations.

**Figure S4:**
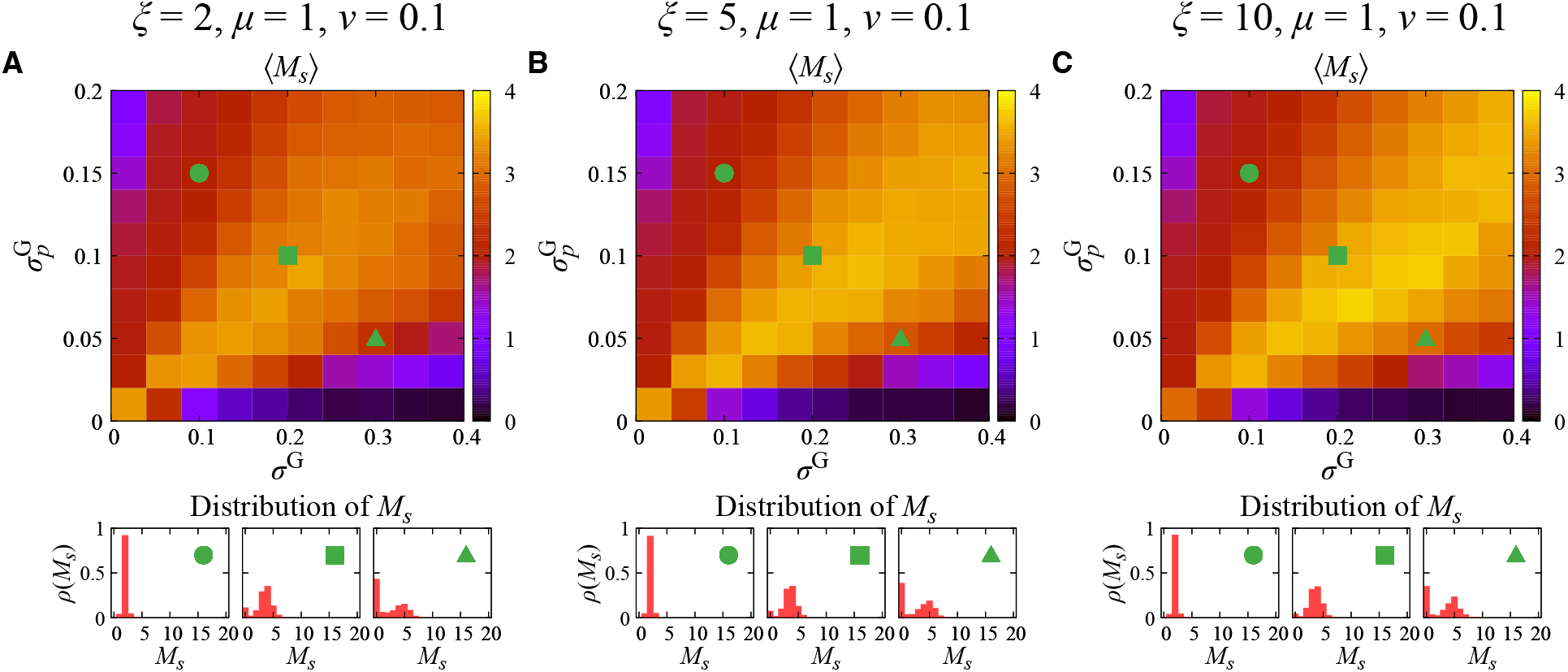
Robustness of phage diversity to the phage burst-size parameter. Average number of surviving phage species ⟨*M*_*s*_⟩ and distributions of *M*_*s*_ for (**A**) *ξ* = 2, (**B**) *ξ* = 5, and (**C**) *ξ* = 10, with *μ* = 1 and *v* = 0.1. All panels use *N* = *M* = 10, Gaussian trait distributions, and the Gaussian infection kernel. The horizontal and vertical axes show 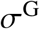 and 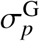, respectively. The lower histograms show the distributions of *M*_*s*_ at the representative parameter sets indicated by the green symbols. Results were averaged over 10^3^ independent trait realizations.

**Figure S5:**
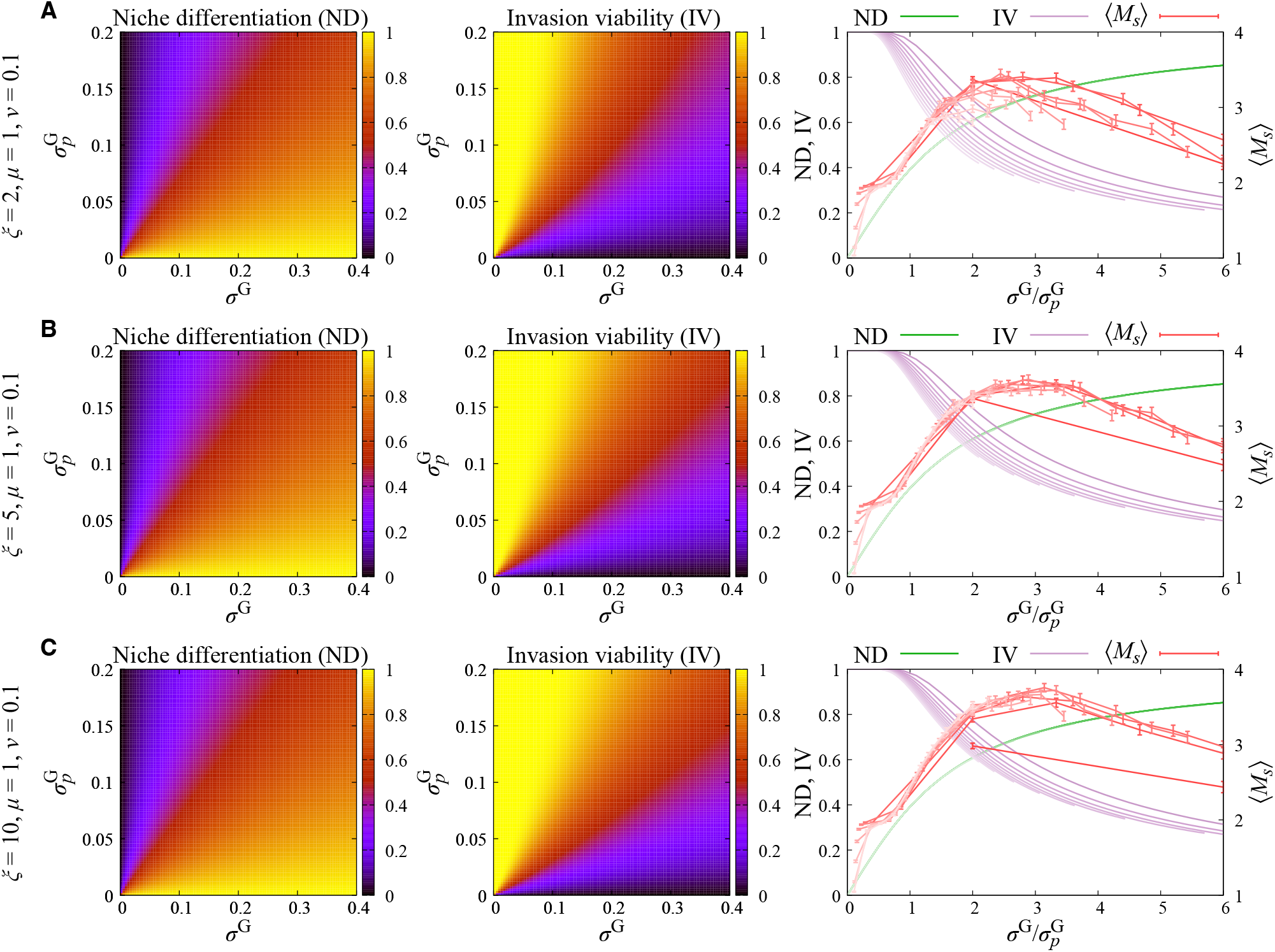
Niche differentiation and invasion viability across phage burst sizes. ND, IV, and ⟨*M*_*s*_⟩ for (**A**) *ξ* = 2, (**B**) *ξ* = 5, and (**C**) *ξ* = 10, with *μ* = 1 and *v* = 0.1. In each row, the left and middle panels show ND and IV in the 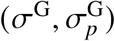 parameter space, and the right panel shows ND, IV, and ⟨*M*_*s*_⟩ as functions of 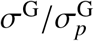. ND is independent of *ξ*, whereas increasing *ξ* relaxes the invasion threshold and increases IV. Each curve corresponds to a fixed 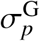 from 0.01 to 0.19 in increments of 0.02. Error bars for ⟨*M*_*s*_⟩ denote standard errors across independent trait realizations.

**Figure S6:**
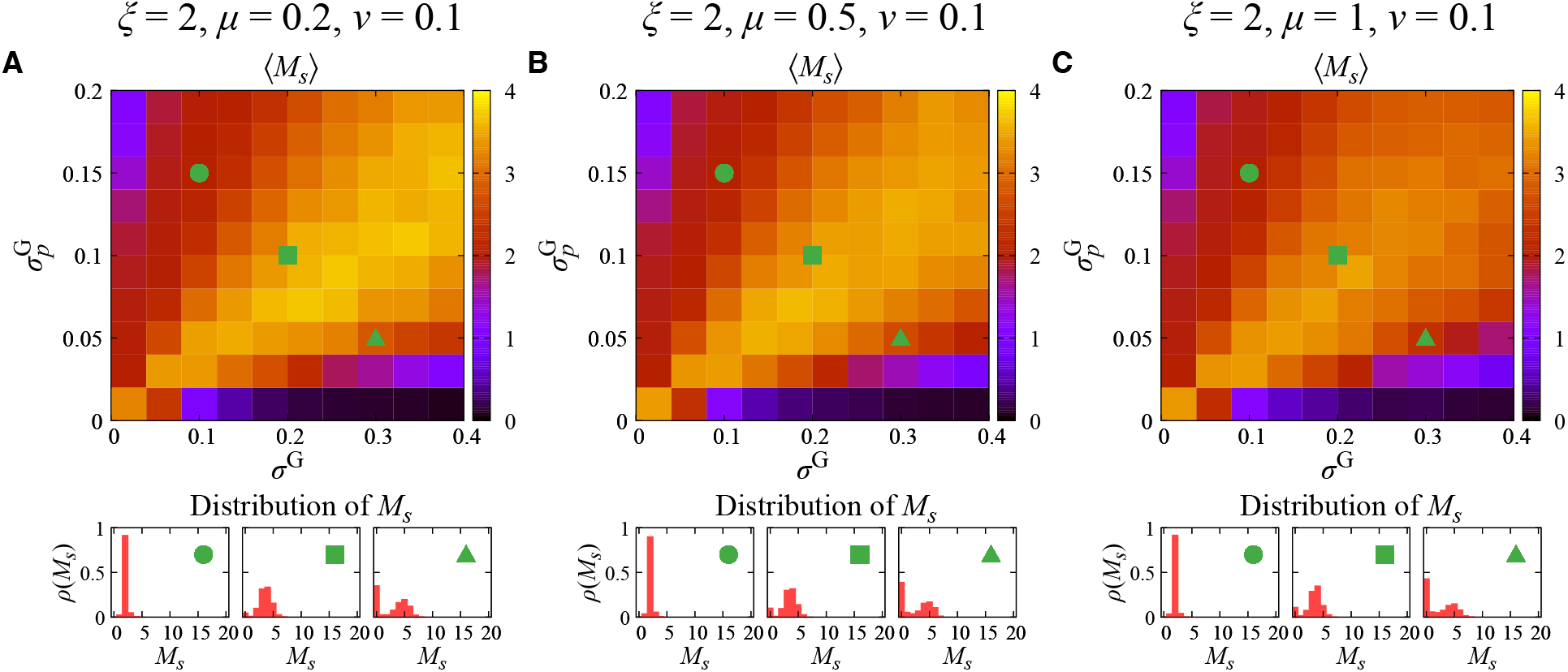
Robustness of phage diversity to the phage degradation rate. Average number of surviving phage species ⟨*M*_*s*_⟩ and distributions of *M*_*s*_ for (**A**) *μ* = 0.2, (**B**) *μ* = 0.5, and (**C**) *μ* = 1, with *ξ* = 2 and *v* = 0.1. All panels use *N* = *M* = 10, Gaussian trait distributions, and the Gaussian infection kernel. The lower histograms show the distributions of *M*_*s*_ at the representative parameter sets indicated by the green symbols. Results were averaged over 10^3^ independent trait realizations.

**Figure S7:**
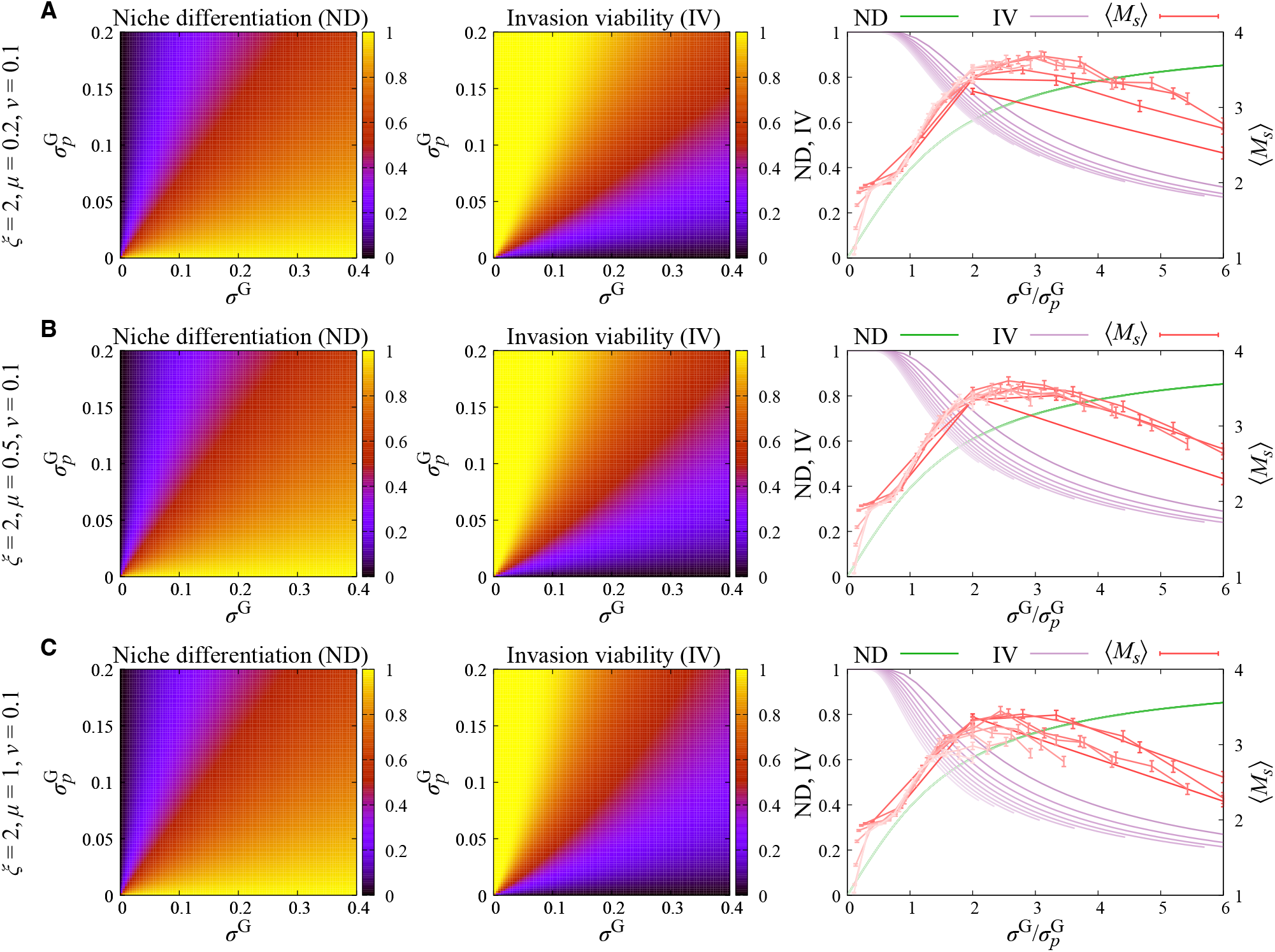
Niche differentiation and invasion viability across phage degradation rates. ND, IV, and ⟨*M*_*s*_⟩ for (**A**) *μ* = 0.2, (**B**) *μ* = 0.5, and (**C**) *μ* = 1, with *ξ* = 2 and *v* = 0.1. In each row, the left and middle panels show ND and IV, and the right panel shows ND, IV, and ⟨*M*_*s*_⟩ as functions of 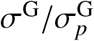. ND is independent of *μ*, whereas increasing *μ* makes the phage invasion condition more restrictive and reduces IV. Each curve corresponds to a fixed 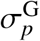 from 0.01 to 0.19 in increments of 0.02. Error bars for ⟨*M*_*s*_⟩ denote standard errors across independent trait realizations.

**Figure S8:**
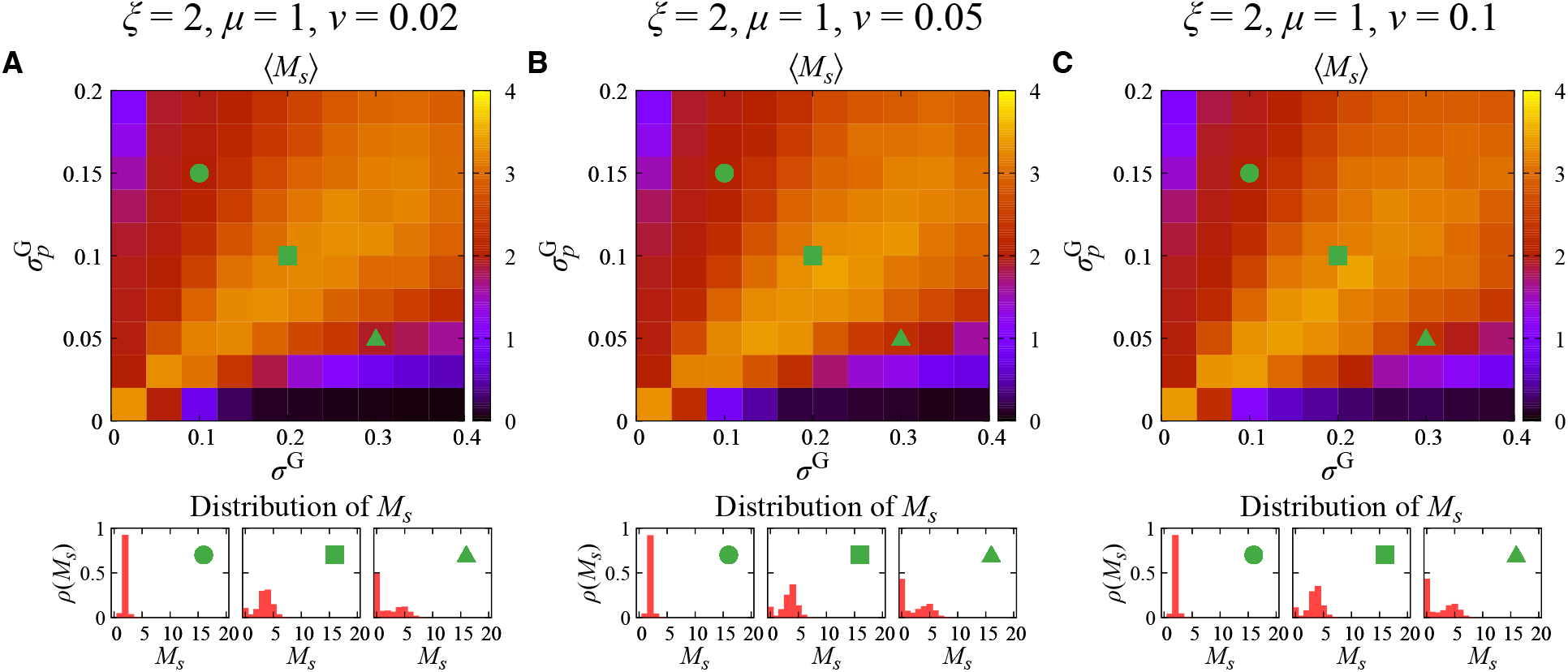
Robustness of phage diversity to the bacterial dilution or mortality rate. Average number of surviving phage species ⟨*M*_*s*_⟩ and distributions of *M*_*s*_ for (**A**) *v* = 0.02, (**B**) *v* = 0.05, and (**C**) *v* = 0.1, with *ξ* = 2 and *μ* = 1. All panels use *N* = *M* = 10, Gaussian trait distributions, and the Gaussian infection kernel. The lower histograms show the distributions of *M*_*s*_ at the representative parameter sets indicated by the green symbols. Results were averaged over 10^3^ independent trait realizations.

**Figure S9:**
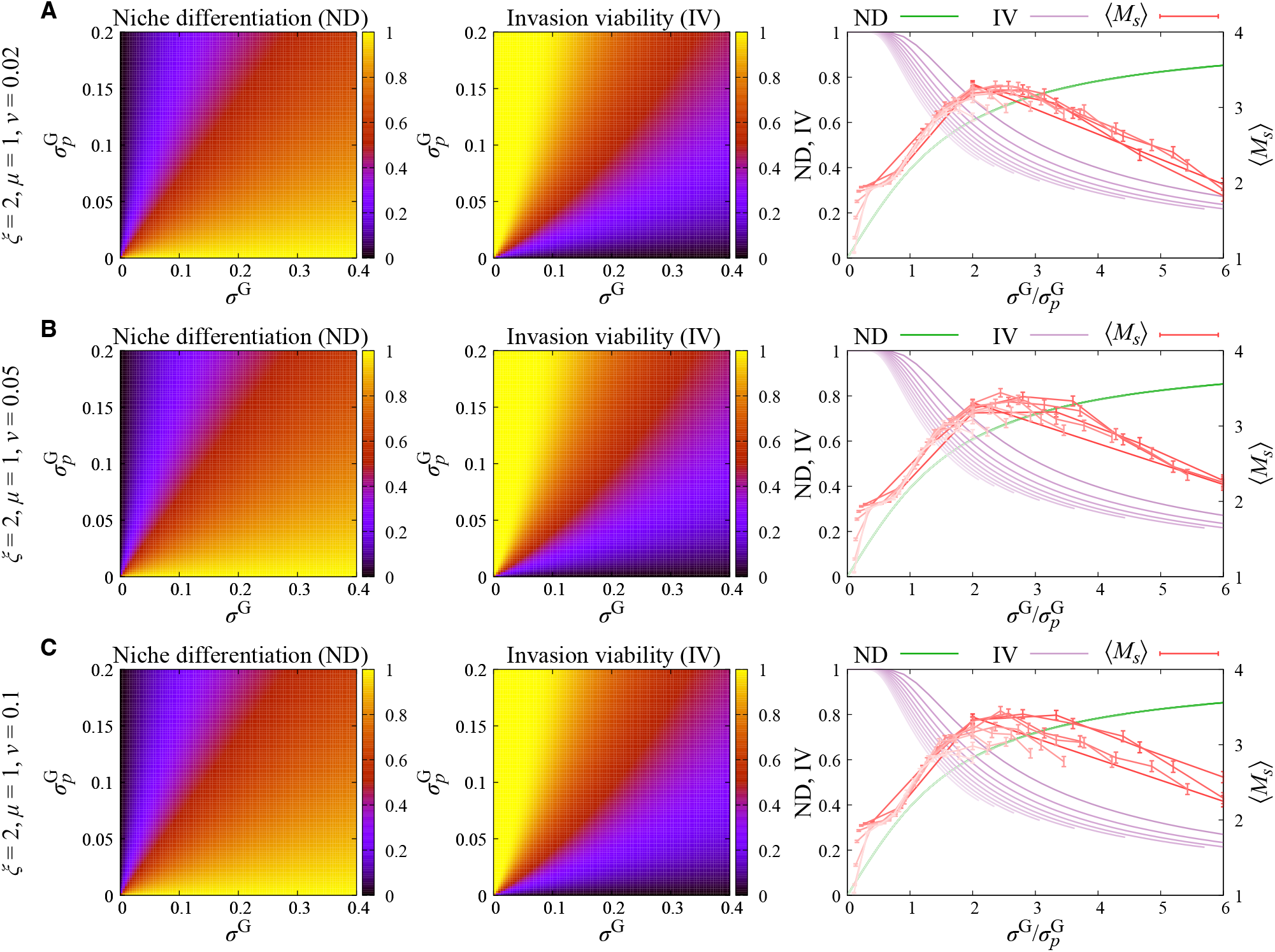
Niche differentiation and invasion viability across bacterial dilution or mortality rates. ND, IV, and ⟨*M*_*s*_⟩ for (**A**) *v* = 0.02, (**B**) *v* = 0.05, and (**C**) *v* = 0.1, with *ξ* = 2 and *μ* = 1. In each row, the left and middle panels show ND and IV, and the right panel shows ND, IV, and ⟨*M*_*s*_⟩ as functions of 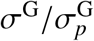. ND is independent of *v*, whereas increasing *v* decreases the bacterial biomass available in the pairwise invasion setting and therefore reduces IV. Each curve corresponds to a fixed 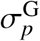 from 0.01 to 0.19 in increments of 0.02. Error bars for ⟨*M*_*s*_⟩ denote standard errors across independent trait realizations.

**Figure S10:**
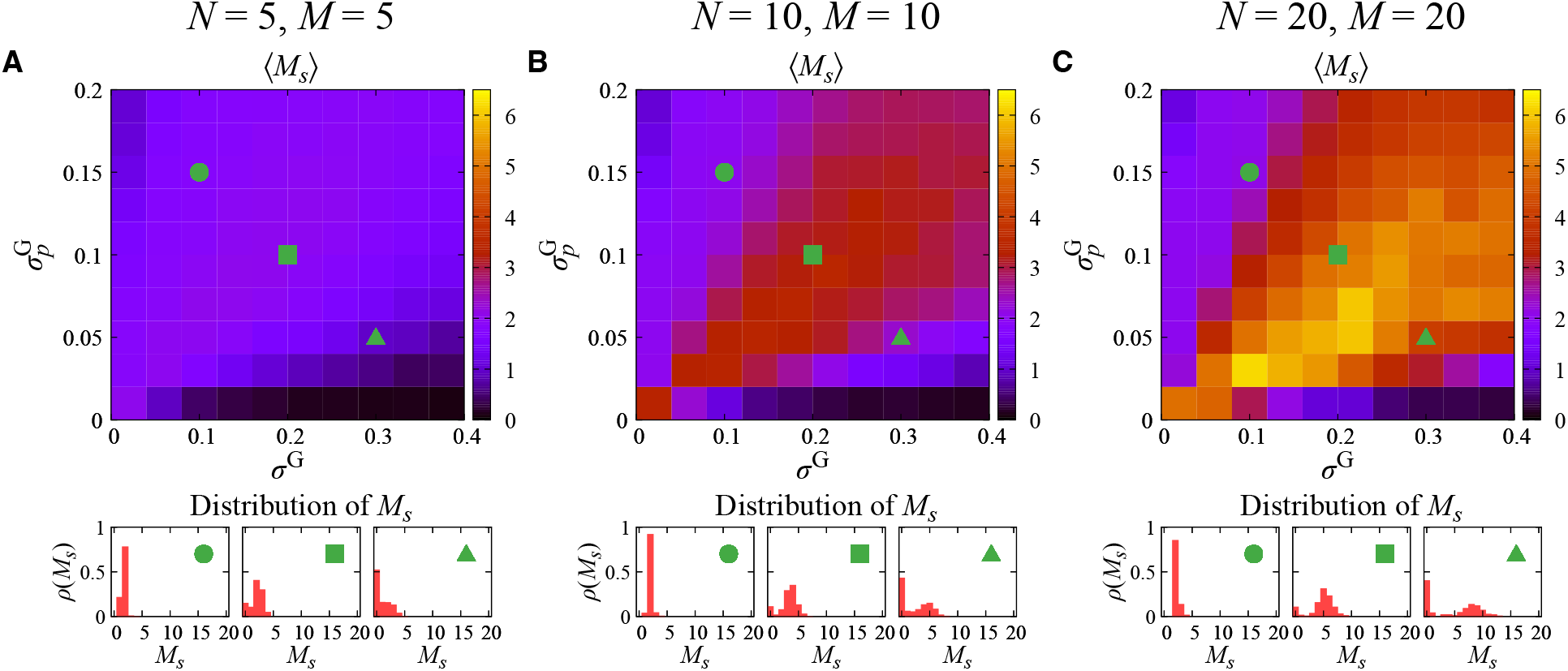
Robustness of phage diversity across community sizes. Average phage richness ⟨*M*_*s*_⟩ and distributions of *M*_*s*_ for Gaussian infection kernels and Gaussian trait distributions with (**A**) *N* = *M* = 5, (**B**) *N* = *M* = 10, and (**C**) *N* = *M* = 20. The horizontal and vertical axes show *σ*^G^ and 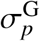, respectively. The lower histograms show the distributions of *M*_*s*_ at the three representative parameter sets indicated by the green symbols. Results were averaged over 10^3^ independent trait realizations for *N* = *M* = 5 and *N* = *M* = 10, and over 2 × 10^2^ realizations for *N* = *M* = 20.

**Figure S11:**
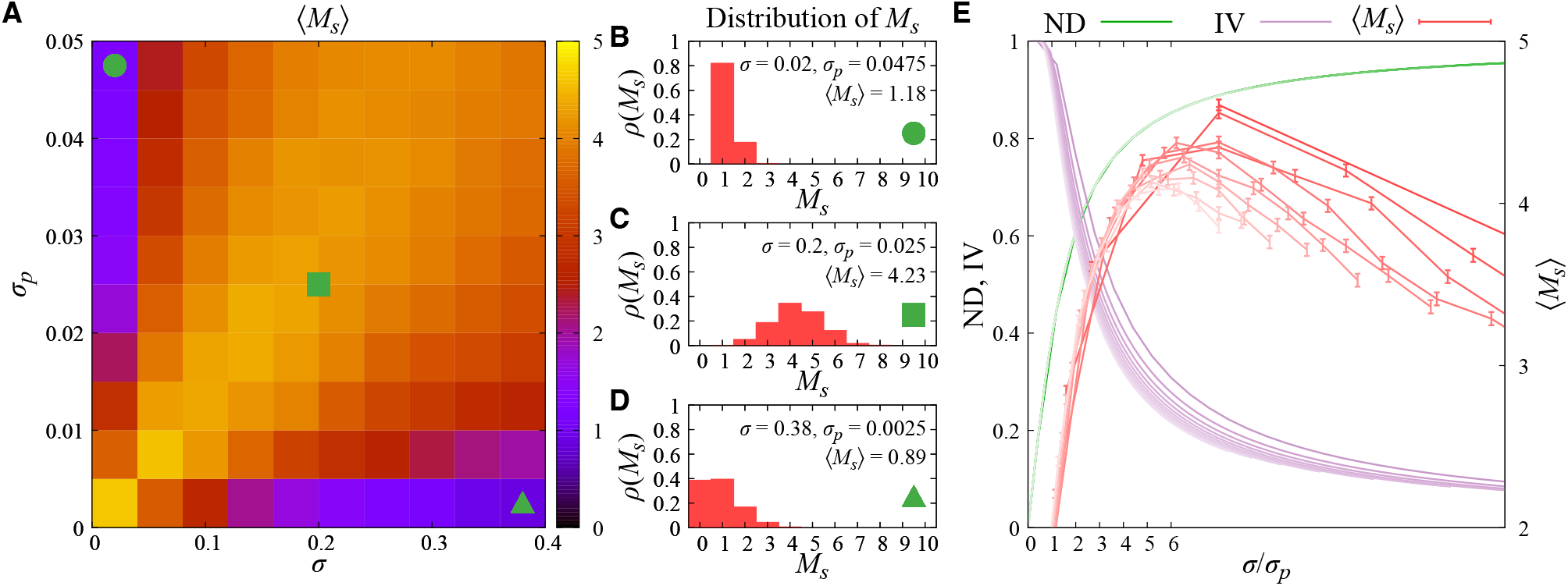
Robustness of the optimal coexistence window to random phenotypic reassignment at bacterial reproduction. In this alternative model, the phenotype of each newly produced bacterial cell is assigned uniformly at random among the *N* predefined phenotypic states, corresponding to complete loss of phenotypic memory at reproduction. (**A**) Average number of surviving phage species ⟨*M*_*s*_⟩ as a function of trait dispersion *σ* and Gaussian infection-kernel width *σ*_*p*_ for *N* = *M* = 10. (**B**–**D**) Distributions of *M*_*s*_ for representative parameter sets corresponding to low trait dispersion and broad infection width (**B**), intermediate scale matching (**C**), and high trait dispersion and narrow infection width (**D**), indicated by the corresponding symbols in (**A**). (**E**) ND, IV, and ⟨*M*_*s*_⟩ plotted as functions of the scale ratio *σ*/*σ*_*p*_. Error bars for ⟨*M*_*s*_⟩ denote standard errors across independent trait realizations. Results were averaged over 10^3^ independent trait realizations.

**Figure S12:**
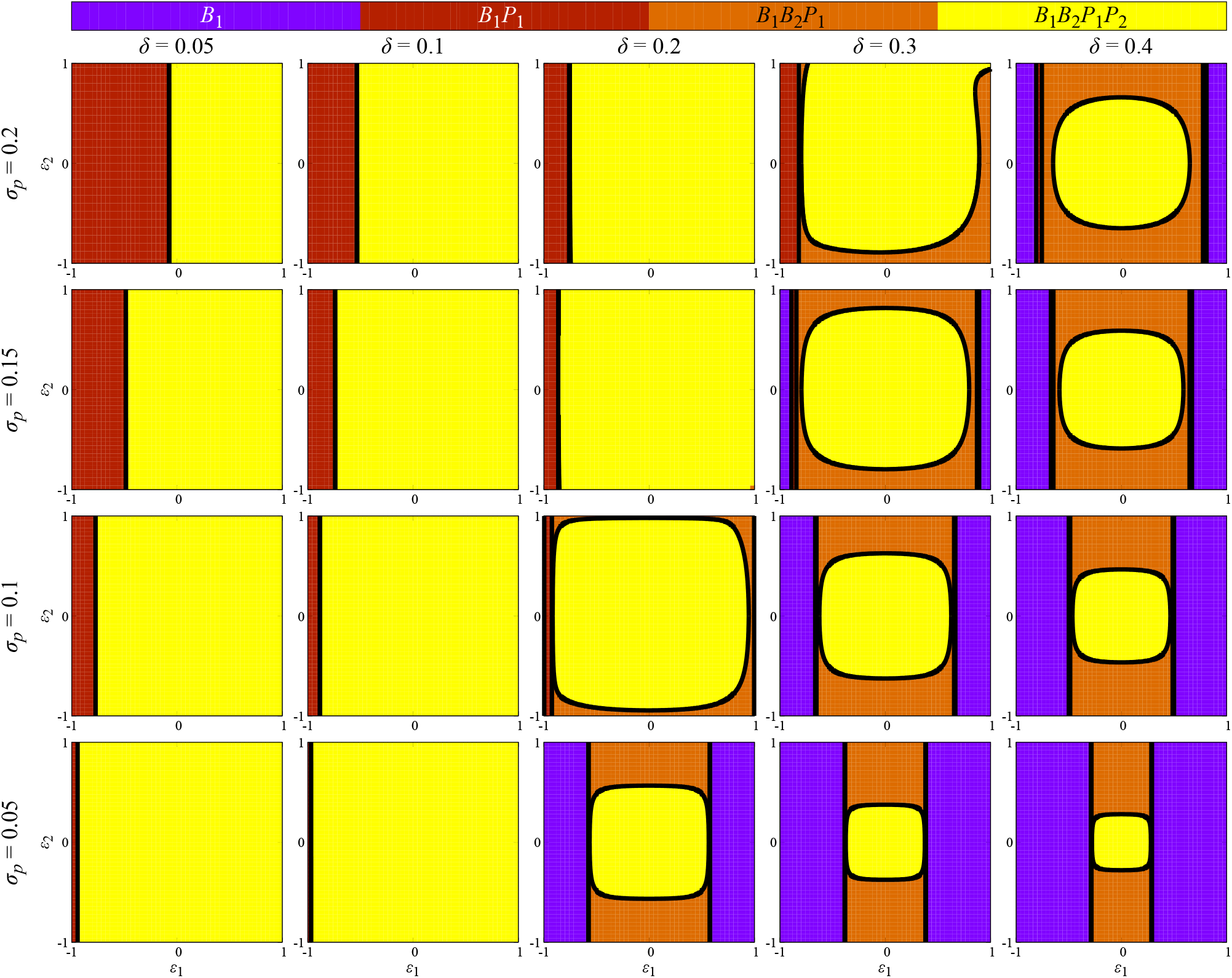
Equilibrium phase diagrams of the minimal two-host–two-phage model. Phases in the (*ϵ*_1_, *ϵ*_2_) plane for different host-state separations *δ* and Gaussian infection-kernel widths *σ*_*p*_. Columns correspond to *δ* = 0.05, 0.1, 0.2, 0.3, and 0.4, and rows correspond to *σ*_*p*_ = 0.20, 0.15, 0.10, and 0.05. Colors indicate the surviving populations: *B*_1_ (purple), *B*_1_*P*_1_ (brown), *B*_1_*B*_2_*P*_1_ (orange), and *B*_1_*B*_2_*P*_1_*P*_2_ (yellow). Solid black curves show the phase boundaries obtained from Eqs. (S61), (S66), and (S75). The full-coexistence region is small when the bacterial states are insufficiently separated, expands at intermediate scale matching, and contracts when their separation becomes too large relative to the infection-kernel width.

**Figure S13:**
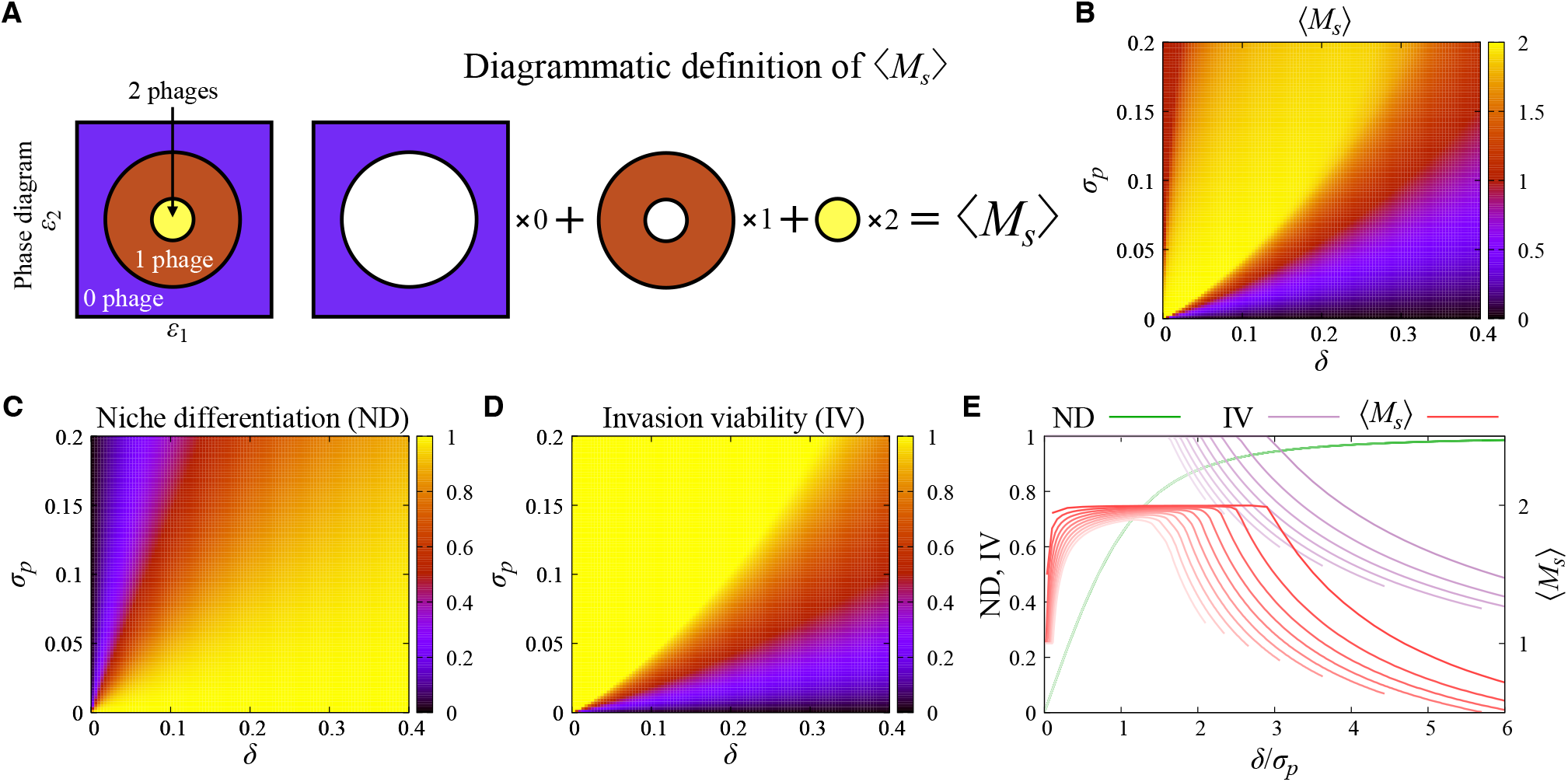
Optimal coexistence in the minimal two-host–two-phage model. (**A**) Diagrammatic definition of ⟨*M*_*s*_⟩ as the area-weighted average of phase-diagram regions with zero, one, and two surviving phages. (**B**) Average phage richness ⟨*M*_*s*_⟩ as a function of *δ* and *σ*_*p*_. (**C**) Niche differentiation ND_2B2P_. (**D**) Invasion viability IV_2B2P_, evaluated using only the matched host–phage pairs (*B*_1_, *P*_1_) and (*B*_2_, *P*_2_). (**E**) ND, IV, and ⟨*M*_*s*_⟩ as functions of the scale ratio *δ*/*σ*_*p*_. ND increases with relative host-state separation, whereas IV decreases at large *δ*/*σ*_*p*_, producing maximal phage coexistence at intermediate scale matching.

## References and Notes

1. R. M. May, Will a Large Complex System be Stable? Nature 238 (5364), 413–414 (1972), doi:10.1038/238413a0, https://doi.org/10.1038/238413a0.

2. T. Thingstad, R. Lignell, Theoretical models for the control of bacterial growth rate, abundance, diversity and carbon demand. Aquatic Microbial Ecology 13, 19–27 (1997), https://www.int-res.com/journals/ame/articles/ame013019.

3. T. F. Thingstad, Elements of a theory for the mechanisms controlling abundance, diversity, and biogeochemical role of lytic bacterial viruses in aquatic systems. Limnology and Oceanography 45 (6), 1320–1328 (2000), doi:10.4319/lo.2000.45.6.1320, https://aslopubs.onlinelibrary.wiley.com/doi/abs/10.4319/lo.2000.45.6.1320.

4. N. Mitarai, A. Marantos, K. Sneppen, Sustainable diversity of phage-bacteria systems. Current Opinion in Systems Biology 35, 100468 (2023), doi:10.1016/j.coisb.2023.100468, https://www.sciencedirect.com/science/article/pii/S2452310023000252.

5. G. F. Gause, The Struggle for Existence. Soil Science 41 (2) (1936), https://journals.lww.com/soilsci/fulltext/1936/02000/the_struggle_for_existence.18.aspx.

6. G. Hardin, The Competitive Exclusion Principle. Science 131 (3409), 1292–1297 (1960), doi:10.1126/science.131.3409.1292, https://www.science.org/doi/abs/10.1126/science.131.3409.1292.

7. S. A. Levin, Community Equilibria and Stability, and an Extension of the Competitive Exclusion Principle. The American Naturalist 104 (939), 413–423 (1970), doi:10.1086/282676, https://doi.org/10.1086/282676.

8. R. D. Holt, Bringing the Hutchinsonian niche into the 21st century: Ecological and evolutionary perspectives. Proceedings of the National Academy of Sciences 106 (supplement 2), 19659–19665 (2009), doi:10.1073/pnas.0905137106, https://www.pnas.org/doi/abs/10.1073/pnas.0905137106.

9. P. A. de Jonge, F. L. Nobrega, S. J. J. Brouns, B. E. Dutilh, Molecular and Evolutionary Determinants of Bacteriophage Host Range. Trends in Microbiology 27 (1), 51–63 (2019), doi:10.1016/j.tim.2018.08.006, https://doi.org/10.1016/j.tim.2018.08.006.

10. S. Heilmann, K. Sneppen, S. Krishna, Coexistence of phage and bacteria on the boundary of self-organized refuges. Proceedings of the National Academy of Sciences 109 (31), 12828–12833 (2012), doi:10.1073/pnas.1200771109, https://www.pnas.org/doi/abs/10.1073/pnas.1200771109.

11. N. Mitarai, S. Brown, K. Sneppen, Population Dynamics of Phage and Bacteria in Spatially Structured Habitats Using Phage λ and Escherichia coli. Journal of Bacteriology 198 (12), 1783–1793 (2016), doi:10.1128/jb.00965-15, https://journals.asm.org/doi/abs/10.1128/jb.00965-15.

12. B. P. Taylor, C. J. Penington, J. S. Weitz, Emergence of increased frequency and severity of multiple infections by viruses due to spatial clustering of hosts. Physical Biology 13 (6), 066014 (2017), doi:10.1088/1478-3975/13/6/066014, https://doi.org/10.1088/1478-3975/13/6/066014.

13. J. J. Bull, et al., Phage-Bacterial Dynamics with Spatial Structure: Self Organization around Phage Sinks Can Promote Increased Cell Densities. Antibiotics 7 (1) (2018), doi:10.3390/antibiotics7010008, https://www.mdpi.com/2079-6382/7/1/8.

14. S. E. Forde, J. N. Thompson, R. D. Holt, B. J. M. Bohannan, COEVOLUTION DRIVES TEMPORAL CHANGES IN FITNESS AND DIVERSITY ACROSS ENVI-RONMENTS IN A BACTERIA–BACTERIOPHAGE INTERACTION. Evolution 62 (8), 1830–1839 (2008), doi:10.1111/j.1558-5646.2008.00411.x, https://doi.org/10.1111/j.1558-5646.2008.00411.x.

15. E. Voigt, B. C. Rall, A. Chatzinotas, U. Brose, B. Rosenbaum, Phage strategies facilitate bacterial coexistence under environmental variability. PeerJ 9, e12194 (2021), doi:10.7717/peerj.12194, https://doi.org/10.7717/peerj.12194.

16. S. J. Beckett, H. T. P. Williams, Coevolutionary diversification creates nested-modular structure in phage–bacteria interaction networks. Interface Focus 3 (6), 20130033 (2013), doi:10.1098/rsfs.2013.0033, https://doi.org/10.1098/rsfs.2013.0033.

17. J. O. Haerter, N. Mitarai, K. Sneppen, Phage and bacteria support mutual diversity in a narrowing staircase of coexistence. The ISME Journal 8 (11), 2317–2326 (2014), doi:10.1038/ismej.2014.80, https://doi.org/10.1038/ismej.2014.80.

18. J. M. Borin, et al., Rapid bacteria-phage coevolution drives the emergence of multiscale networks. Science 382 (6671), 674–678 (2023), doi:10.1126/science.adi5536, https://www.science.org/doi/abs/10.1126/science.adi5536.

19. N. C. Pyenson, et al., Diverse phage communities are maintained stably on a clonal bacterial host. Science 386 (6727), 1294–1300 (2024), doi:10.1126/science.adk1183, https://www.science.org/doi/abs/10.1126/science.adk1183.

20. D. I. Bolnick, et al., Why intraspecific trait variation matters in community ecology. Trends in Ecology & Evolution 26 (4), 183–192 (2011), doi:10.1016/j.tree.2011.01.009, https://doi.org/10.1016/j.tree.2011.01.009.

21. S. Des Roches, et al., The ecological importance of intraspecific variation. Nature Ecology & Evolution 2 (1), 57–64 (2018), doi:10.1038/s41559-017-0402-5, https://doi.org/10.1038/s41559-017-0402-5.

22. S. M. Stump, C. Song, S. Saavedra, J. M. Levine, D. A. Vasseur, Synthesizing the effects of individual-level variation on coexistence. Ecological Monographs 92 (1), e01493 (2022), doi: 10.1002/ecm.1493, https://esajournals.onlinelibrary.wiley.com/doi/abs/10.1002/ecm.1493.

23. A. J. Lotka, Elements of physical biology (Williams & Wilkins) (1925).

24. V. Volterra, Variazioni e fluttuazioni del numero d’individui in specie animali conviventi (Mem. Acad. Lincei Roma) (1926).

25. A. Campbell, Conditions for the Existence of Bacteriophage. Evolution 15 (2), 153–165 (1961), http://www.jstor.org/stable/2406076.

26. L. F. Jover, M. H. Cortez, J. S. Weitz, Mechanisms of multi-strain coexistence in host–phage systems with nested infection networks. Journal of Theoretical Biology 332, 65–77 (2013), doi: 10.1016/j.jtbi.2013.04.011, https://www.sciencedirect.com/science/article/pii/S0022519313001641.

27. L. F. Jover, C. O. Flores, M. H. Cortez, J. S. Weitz, Multiple regimes of robust patterns between network structure and biodiversity. Scientific Reports 5 (1), 17856 (2015), doi:10.1038/srep17856, https://doi.org/10.1038/srep17856.

28. A. Marantos, N. Mitarai, K. Sneppen, From kill the winner to eliminate the winner in open phage-bacteria systems. PLOS Computational Biology 18 (8), 1–15 (2022), doi: 10.1371/journal.pcbi.1010400, https://doi.org/10.1371/journal.pcbi.1010400.

29. R. MacArthur, R. Levins, The Limiting Similarity, Convergence, and Divergence of Coexisting Species. The American Naturalist 101 (921), 377–385 (1967), doi:10.1086/282505, https://www.journals.uchicago.edu/doi/abs/10.1086/282505.

30. J. S. Weitz, H. Hartman, S. A. Levin, Coevolutionary arms races between bacteria and bacteriophage. Proceedings of the National Academy of Sciences 102 (27), 9535–9540 (2005), doi:10.1073/pnas.0504062102, https://www.pnas.org/doi/abs/10.1073/pnas.0504062102.

31. H. T. Williams, Phage-induced diversification improves host evolvability. BMC Evolutionary Biology 13 (1), 17 (2013), doi:10.1186/1471-2148-13-17, https://doi.org/10.1186/1471-2148-13-17.

32. G. A. Cooper, M. Liu, J. Peña, S. A. West, The evolution of mechanisms to produce phenotypic heterogeneity in microorganisms. Nature communications 13 (1), 195 (2022), https://www.nature.com/articles/s41467-021-27902-4.

33. X. Liang, et al., Bacteriophage-driven microbial phenotypic heterogeneity: ecological and biogeochemical importance. npj Biofilms and Microbiomes 11 (1), 82 (2025), https://www.nature.com/articles/s41522-025-00727-5.

34. M. Zhu, X. Dai, Shaping of microbial phenotypes by trade-offs. Nature Communications 15 (1), 4238 (2024), https://www.nature.com/articles/s41467-024-48591-9.

35. M. Ackermann, A functional perspective on phenotypic heterogeneity in microorganisms. Nature Reviews Microbiology 13 (8), 497–508 (2015), doi:10.1038/nrmicro3491, https://doi.org/10.1038/nrmicro3491.

36. Z. Bódi, et al., Phenotypic heterogeneity promotes adaptive evolution. PLoS biology 15 (5), e2000644 (2017).

37. M. Amicone, I. Gordo, Molecular signatures of resource competition: Clonal interference favors ecological diversification and can lead to incipient speciation. Evolution 75 (11), 2641–2657 (2021).

38. C. O. Flores, J. R. Meyer, S. Valverde, L. Farr, J. S. Weitz, Statistical structure of host–phage interactions. Proceedings of the National Academy of Sciences 108 (28), E288– E297 (2011), doi:10.1073/pnas.1101595108, https://www.pnas.org/doi/abs/10.1073/pnas.1101595108.

39. C. O. Flores, S. Valverde, J. S. Weitz, Multiscale structure and geographic drivers of cross-infection within marine bacteria and phages. The ISME Journal 7 (3), 520–532 (2013), doi: 10.1038/ismej.2012.135, https://doi.org/10.1038/ismej.2012.135.

40. J. S. Weitz, et al., Phage–bacteria infection networks. Trends in Microbiology 21 (2), 82–91 (2013), doi:10.1016/j.tim.2012.11.003, https://doi.org/10.1016/j.tim.2012.11.003.

41. D. A. Korytowski, H. L. Smith, How nested and monogamous infection networks in host-phage communities come to be. Theoretical Ecology 8 (1), 111–120 (2015), doi:10.1007/s12080-014-0236-6, https://doi.org/10.1007/s12080-014-0236-6.

42. J. I. Park, D.-S. Lee, S. H. Lee, H. J. Park, Incorporating Heterogeneous Interactions for Ecological Biodiversity. Phys. Rev. Lett. 133, 198402 (2024), doi:10.1103/PhysRevLett.133.198402, https://link.aps.org/doi/10.1103/PhysRevLett.133.198402.

43. C. Xue, N. Goldenfeld, Coevolution Maintains Diversity in the Stochastic “Kill the Winner” Model. Phys. Rev. Lett. 119, 268101 (2017), doi:10.1103/PhysRevLett.119.268101, https://link.aps.org/doi/10.1103/PhysRevLett.119.268101.

44. O. Kimchi, Y. Meir, N. S. Wingreen, Bacterial defense and phage counterdefense lead to coexistence in a modeled ecosystem. Proceedings of the National Academy of Sciences 121 (44), e2414229121 (2024), doi:10.1073/pnas.2414229121, https://www.pnas.org/doi/abs/10.1073/pnas.2414229121.

45. W. Cui, J. M. Fendley, S. Srikant, B. I. Shraiman, A minimal model of panimmunity maintenance by horizontal gene transfer in the ecological dynamics of bacteria and phages. Proceedings of the National Academy of Sciences 122 (31), e2417628122 (2025), doi:10.1073/pnas.2417628122, https://www.pnas.org/doi/abs/10.1073/pnas.2417628122.

46. K. Biswas, N. Brenner, Universality of phenotypic distributions in bacteria. Physical Review Research 6 (2), L022043 (2024).

47. GitHub, https://github.com/JungHoKim4823/PhageDiversity2026.

